# ALS-Associated TDP-43 Dysfunction Compromises UPF1-Dependent mRNA Metabolism Pathways Including Alternative Polyadenylation and 3’UTR Length

**DOI:** 10.1101/2024.01.31.578311

**Authors:** Francesco Alessandrini, Matthew Wright, Tatsuaki Kurosaki, Lynne E. Maquat, Evangelos Kiskinis

## Abstract

UPF1-mediated decay entails several mRNA surveillance pathways that play a crucial role in cellular homeostasis. However, the precise role of UPF1 in postmitotic neurons remains unresolved, as does its activity in amyotrophic lateral sclerosis (ALS), a devastating neurodegenerative disease characterized by TDP-43 pathology and disrupted mRNA metabolism. Here, we used human iPSC-derived spinal motor neurons (MNs) to identify mRNAs subject to UPF1 degradation by integrating RNA-seq before and after UPF1 knockdown with RIP-seq to identify RNAs that co-immunoprecipitate with the active form of phosphorylated UPF1. We define a stringent set of *bona fide* UPF1 targets in MNs that are functionally enriched for autophagy and structurally enriched for GC-rich and long 3’ UTRs but not for premature termination codon (PTC)-containing transcripts. TDP-43 depletion in iPSC-derived MNs reduces UPF1 phosphorylation and consequently post-transcriptional upregulation of UPF1 targets, suggesting that TDP-43 dysfunction compromises UPF1-mediated mRNA surveillance. Intriguingly, our datasets reveal that UPF1 and TDP-43 regulate alternative polyadenylation and 3’UTR length of mRNAs associated with synaptic and axonal function, a process that we find to be compromised in ALS models *in vitro* and ALS patient tissue. Our study provides a comprehensive description of UPF1-mediated mRNA decay activity in neurons, reveals overlapping roles between UPF1 and TDP-43 in regulating 3’UTR length, and offers novel insight into the intricate interplay between RNA metabolism and neurodegeneration in ALS.

## INTRODUCTION

Amyotrophic Lateral Sclerosis (ALS) is a devastating neurodegenerative disease stemming from the destruction of motor neurons (MNs) that reside in the brain and spinal cord (Goutman et al., 2022; Rowland and Shneider, 2001). Its relentless progression eventually leads to a complete loss of motor function, and results in fatality within 2-5 years after initial diagnosis. More than 30 genes have been identified as causative or disease-modifying, with pathogenic variants in *C9ORF72, SOD1, TARDBP, FUS* and *NEK1* occurring most frequently (Akcimen et al., 2023; Suzuki et al., 2023; Taylor et al., 2016). However, the underlying etiology of ALS remains elusive, and treatment options for the majority of sporadic patients remain limited. The clearance of TAR DNA-binding protein 43 (TDP-43) from the nucleus and its concordant accumulation in ubiquitinated and hyperphosphorylated cytosolic aggregates is a neuropathological hallmark of most ALS patients (Neumann et al., 2006; Vatsavayai et al., 2016), as it is observed in approximately 97% of all patients regardless of their genetic predisposition (Tan et al., 2017).

TDP-43 is a multi-faceted RNA/DNA-binding protein that can shuttle between the nucleus and cytoplasm but under physiological conditions is localized predominantly in the nucleus (Chen-Plotkin et al., 2010). It acts as a master regulator of various aspects of mRNA metabolism including trafficking and post-transcriptional splicing (Ayala et al., 2011; Fiesel et al., 2012; Klim et al., 2021 Swain, 2016 #51; Ling et al., 2015; Modic et al., 2019; Polymenidou et al., 2011; Rot et al., 2017). While the gain-of-function effects associated with TDP-43 aggregation have been a major focus since the initial identification of TDP-43-positive inclusions in ALS patients, there has been renewed interest recently in understanding how TDP-43 loss-of-function impacts RNA splicing (Akiyama et al., 2022; Klim et al., 2021; Mehta et al., 2023). Studies utilizing neurons derived from induced pluripotent stem cells (iPSCs) as well as postmortem ALS patient brain tissue have catalogued a series of TDP-43 dependent mis-splicing events in genes such as *STMN2* and *UNC13A* (Brown et al., 2022; Klim et al., 2019; Ma et al., 2022; Melamed et al., 2019). Some of these novel mRNA species include cryptic or skiptic exons, which can give rise to neopeptides (Seddighi et al., 2023), or premature termination codons (PTCs) that earmark them for degradation (Brown et al., 2022; Ma et al., 2022).

The integrity and quality of newly synthesized mRNAs are supervised by a number of post-transcriptional surveillance mechanisms (Wolin and Maquat, 2019). Up-frameshift protein 1, encoded by the *UPF1* gene, is a multifunctional enzyme retaining RNA-dependent helicase and ATPase activity that plays a critical role in several mRNA quality control mechanisms, including nonsense-mediated decay (NMD) (Hwang et al., 2021; Karousis et al., 2016; Kurosaki et al., 2021a; Lejeune et al., 2003; Popp and Maquat, 2018), Staufen1 mediated decay (SMD) (Park and Maquat, 2013), structure-mediated decay (SRD) (Fischer et al., 2020), histone mRNA decay (HMD) (Kaygun and Marzluff, 2005), and TSN-mediated microRNA decay (TumiD) (Elbarbary et al., 2017). UPF1 operates within a collective framework with other proteins such as eRF1, UPF2 and UPF3X, as well as the kinases SMG1 and AKT that is responsible for its phosphorylation and activation (Kurosaki et al., 2019). During the pioneer round of translation, phosphorylated UPF1 recruits degradative activities that selectively degrade mRNAs harboring a PTC situated upstream of an exon-junction complex (Kurosaki et al., 2019). Notably, the dynamic activity of UPF1 and associated proteins is finely regulated through a feedback loop within the degradation pathway (Huang et al., 2011; Yepiskoposyan et al., 2011). Beyond the decay of PTC-encoding transcripts, phosphorylated UPF1 is also known to recruit degradative activities to approximately 5-10% of physiological mRNAs so as to modulate the expression of genes involved in cellular responses to stress stimuli (Kurosaki et al., 2019).

While UPF1 has been extensively studied in dividing cells, the precise role of UPF1-dependent mRNA decay in postmitotic neurons remains unresolved. It has been shown to enable compartmentalized gene expression, modulate synaptic activity and axon guidance, and regulate the expression of neuronal mRNAs that are critical for mouse brain development (Colak et al., 2013; Kurosaki et al., 2021b; Tan et al., 2020). Given the extent of defective mRNA metabolism in ALS, it is unsurprising that alterations in UPF1-mediated NMD activity have been highlighted in several studies. We and others have shown that overexpression of UPF1 rescues neurotoxicity in genetic ALS models *in vitro* and *in vivo,* although this mechanism remains controversial. (Barmada et al., 2015; Jackson et al., 2015; Ortega et al., 2020; Zaepfel et al., 2021). Additionally, components of the NMD machinery have been shown to be transcriptionally activated in FUS and C9ORF72 ALS models (Gittings et al., 2023; Kamelgarn et al., 2018; Sun et al., 2020), and NMD has been shown to be dysregulated in single-cell analyses of postmortem ALS patient tissue (Gittings et al., 2023). Thus, whether and how mRNA surveillance, and more specifically UPF1 activity, are impacted by ALS-associated disease pathophysiology remain unknown. What also remains unclear is how the emergence of novel mis-spliced mRNAs downstream of TDP-43 dysfunction engage with UPF1 activity.

The present study focused on defining the role of UPF1 in human MNs, which represent the most vulnerable cellular population in ALS, and understanding the interplay between TDP-43 dysfunction and UPF1-mediated degradation of mRNAs. Using human iPSC-based models, we catalogue a high-confidence list of UPF1 target mRNAs that are functionally enriched for autophagy and structurally enriched for long 3′-untranslated regions (3’UTRs). We find that loss-of-function of TDP-43 impairs the efficiency of UPF1 activity and allows for the buildup of erroneously processed mRNAs. Notably, we make the surprising finding that UPF1 and TDP-43 loss-of-function disrupt alternative polyadenylation (APA) and 3’UTR length of overlapping transcripts, a process that we find to be compromised in neuronal models of ALS *in vitro* and ALS postmortem patient tissue. Our findings resolve long-standing questions regarding the functional role of UPF1 in ALS neurons and highlight a disruption in APA site selection as a critical neuropathological mechanism in ALS that is regulated by the actions of TDP-43 and UPF1.

## RESULTS

### Defining *bona fide* UPF1 Target Transcripts in Human MNs

In order to identify transcripts that are targeted and degraded by UPF1-mediated mechanisms in human motor neurons (MNs), we used an established protocol (Alvarez et al., 2023; Ziller et al., 2018) to differentiate a healthy control iPSC line into spinal MNs (iPSC line 18a) (Boulting et al., 2011). We matured the cultures for 40 days *in vitro* and performed RNA immunoprecipitation (RIP) followed by deep-sequencing (seq) of RNAs bound to the hyper-phosphorylated, i.e. activated, form of UPF1 (pUPF1) (Fig. 1A). To increase the cellular level of pUPF1, we treated the MN cultures with 50 nM of the protein phosphatase inhibitor okadaic acid for 2 hours just prior to cell lysis and immunoprecipitation (Fig. S1A) (Isken et al., 2008; Kurosaki et al., 2018; Kurosaki et al., 2014). RIP-seq analyses identified 5,469 genes that were bound by pUPF1 relative to an IgG control (n=4 independent differentiations, p<0.05, fold change>1.2) (Fig. 1E left). In parallel, we used successive siRNA treatments during the course of 15 days to knockdown 80% of *UPF1* RNA and 70-80% of UPF1 protein (Fig. 1A-C and Fig. S1C), without causing detectable cell death (Fig. S1B). The reduction in UPF1 levels caused the significant upregulation of 1,961 genes measured by whole transcriptome RNA-seq (n=5 independent differentiations, adj. p<0.05, fold change>1.2) (Fig. 1E left and Fig. S1C). To definitively ascribe the upregulation of genes to a reduction of UPF1-dependent posttranscriptional decay we applied REMBRANDTS (<u>rem</u>oving <u>b</u>ias from <u>R</u>NA-seq <u>an</u>alysis of <u>d</u>ifferential transcript <u>s</u>tability), an RNA-seq tool that allows for assessing differential mRNA half-lives (Alkallas et al., 2017). This approach allowed us to estimate in an unbiased fashion the differential abundance of mature mRNA relative to the corresponding pre-mRNA (i.e. the delta (Δ) between exonic and intronic sequences) for every gene in UPF1 knockdown (KD) and scrambled control conditions (Fig. 1A). This analysis identified 1,850 genes with increased mature to premature mRNA levels, that are likely degraded by pUPF1-dependent mechanisms (p<0.05) (Fig. 1E). As expected, *UPF1* mRNA itself was downregulated post-transcriptionally by siRNA treatment, while other transcripts such as *IPO9* were upregulated post-transcriptionally after UPF1 KD, likely due to inhibition of mature mRNA decay (Fig. 1D). Integration of all 3 datasets revealed a cohort of mRNAs associated with 639 genes that met all three criteria, i.e., were bound by pUPF1, were upregulated in expression after UPF1 KD, and exhibited an increased mature mRNA to pre-mRNA ratio (Fig. 1E). These mRNAs represent *bona fide* or high-confidence UPF1-mediated decay targets (referred to as UPF1-targets hereafter).

**Figure 1.**
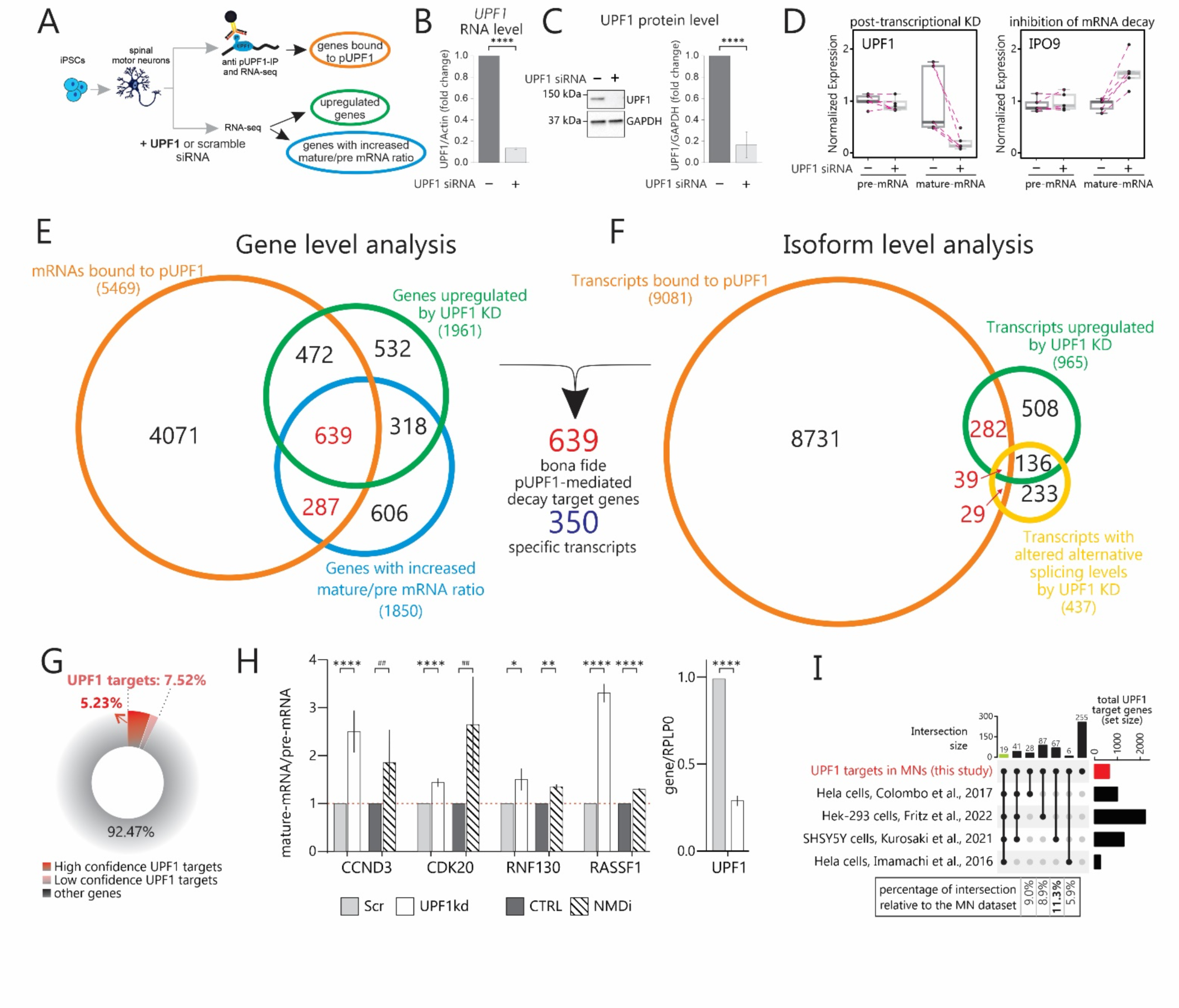
Defining *bona fide* UPF1 Target Transcripts in Human MNs. **A)** Experimental schematic used to identify mRNAs bound to pUPF1 and the post-transcriptomic modulation in response to UPF1 KD in iPSC-derived hMNs. **B)** Bar plot showing the RNA level of *UPF1* in hMNs after siRNA treatment, analyzed by qPCR and normalized to the level of housekeeping gene Actin (p<0.0001). **C)** Left: representative WB for UPF1 in MN cultures after treatment with siRNA targeting *UPF1*. GAPDH was used as a loading control. Right: bar plot showing the fold change in the UPF1/GAPDH ratio after siRNA treatment (t.test, p<0.0001). **D)** Boxplots showing pre-mRNA and mature mRNA modulation of *UPF1* and *IPO9* after treatment with siRNA targeting *UPF1* by RNA-seq data. Pink dotted lines connect samples from the same differentiation (n=5 independent differentiations). **E)** Venn diagram displaying the overlap of mRNAs bound to pUPF1 (n=3 individual differentiations), the genes showing overexpression upon UPF1 KD and genes showing decreased pre-mRNA/mature mRNA ratio upon UPF1 KD (n=5 individual differentiations). **F)** Venn diagram displaying the overlap of specific isoforms bound to pUPF1, the transcripts showing overexpression upon UPF1 KD and transcripts showing differential transcriptional usage upon UPF1 KD. **G)** Donut chart showing the percentage of protein coding genes targeted by UPF1 with high and low confidence relative to the total of the protein coding genes expressed by hMNs. **H)** Left: Bar plot showing the fold change of the pre-mRNA/mature mRNA ratio of selected UPF1-target genes in hMNs treated with scramble or UPF1 siRNA and in control and NMDi treated hMNs, analyzed by qPCR. Right: Right: Bar plot showing the fold change of UPF1 after treatment with siRNA, analyzed by qPCR and normalized to the level of housekeeping gene RPLP0 (3 to 4 independent differentiations, paired t.test, *p<0.05; **p<0.01; ****p<0.0001; ##p<0.001 when t.test performed on technical replicates from 2 independent differentiations). **I)** Upset plot showing selected intersections of UPF1-targets identified in the present study and NMD targets identified in previous studies.

We next utilized Salmon and Fishpond to align reads from RIP-seq and identify specific isoforms rather than genes, that are preferentially bound to pUPF1. We intersected this dataset with isoforms upregulated or exhibiting positive differential transcriptional usage (DTU) after knocking down UPF1 and earmarked 350 transcripts associated with 322 genes that are putatively targeted for UPF1-mediated degradation in human MNs (Fig. 1F). Notably, in some cases, UPF1 modulation resulted in the upregulation of all isoforms associated with a gene, while in other cases only a specific isoform was affected. Combining the datasets from the gene level and isoform level analysis highlighted between 794 and 1093 mRNAs that are modulated by UPF1 with high and low confidence respectively (Fig. S1D). UPF1 targets represent 4.7-6.2% of all genes analyzed in these experiments (Fig. S1E), and 5.2-7.5% when considering exclusively protein coding genes (Fig. 1G). Our findings are consistent with previous studies in non-neuronal cells showing that approximately 5-10% of the protein-coding cellular genes are modulated by NMD (Kurosaki et al., 2019).

Critically, the mRNAs we identified as UPF1 targets were not dependent on genetic background, as we were able to recapitulate the upregulation for the overwhelming majority of them by knocking down UPF1 in day 40 MNs derived from another iPSC line (iPSC line 11a) (Boulting et al., 2011) and performing whole transcriptome RNA-seq (Fig. S1F). To verify the post-transcriptional UPF1-dependent regulation using an alternative method, we designed qRT-PCR primers able to distinguish between pre-mRNA and mature mRNA for a select number of genes. Using this approach, we observed an increase in the mature mRNA/pre-mRNA ratio after *UPF1* KD in MNs derived from several additional independent differentiations (Fig. 1H and Fig. S1G). We observed a similar increase in MN cultures treated with a small molecule inhibitor of NMD (NMDi-1, 0.5 μM, 72hrs) for 11/12 target transcripts analyzed, further suggesting that the regulation is dependent on the activity of UPF1 (Fig. 1H and Fig. S1G). Lastly, to assay for the specificity of these UPF1-dependent events in MNs, we compared the list of our high-confidence UPF1 targets with findings from 4 other recent studies (Fig. 1I and Fig. S1H-I) (Colombo et al., 2017; Fritz et al., 2022; Imamachi et al., 2017; Kurosaki et al., 2021a). Only 19 targets were found to be modulated by UPF1 across all studies, while an additional 60 were shared amongst at least 4 studies. These overlapping targets do not exhibit strong functional interconnectedness (Fig. S1J), but they do exhibit enrichment for biological processes including “positive regulation of transcription from RNA polymerase II promoter in response to stress”, as well as involvement in pathways like “protein processing in the ER”, and “apoptosis”. Among the transcripts we identified in MNs, 60% have been previously associated with NMD in other cells, while 40% represent novel UPF1 targets (Fig. S1H). Notably, these novel UPF1 target genes are functionally enriched within brain tissue and the central nervous system (Fig. S1K-L), suggesting some degree of neuronal specificity.

### UPF1 Target Transcripts in MNs Exhibit Long and Highly Structured 3’UTRs

Having established a list of *bona fide* UPF1 targets in MNs we next sought to define whether they share any common structural or functional properties. We focused on the 350 high-confidence mRNAs plus specific transcripts that represent the most modulated isoform for a gene we classified as a UPF1 target. As expected, the expression of these transcripts showed a significant increase upon UPF1 KD relative to all transcripts detected in our RNA-seq that we used as background for our analysis (Fig. S2A). We first examined how each mRNA and associated gene were annotated by ENSEMBL. The overwhelming majority of transcripts, represented by 89%, were protein-coding, with the remaining annotated as pseudogenes or long non-coding RNAs (Fig. 2A). Among protein-coding genes, two-thirds of the transcripts identified as UPF1 targets were classified as “protein coding”, while the remaining transcripts were annotated as “nonsense-mediated decay”, “processed transcripts” or “retaining intron” sequences (Fig. 2A). We then focused our analysis on characteristics determined by each specific nucleotide sequence, comparing the UPF1 target transcripts to all the other transcripts detected by RNA-seq (referred to as background transcripts hereafter). Nearly all mRNA targets (92.5%) featured at least one predicted pUPF1 GC-rich binding motif (e.g., CCUG[G/A][G/A][G/A]) (Imamachi et al., 2017) within their 3’UTR, with the majority encoding multiple sites and about half harboring more than six sites, which likely contribute to stabilizing the interaction between pUPF1 and mRNA molecules (Fig. 2B-C and Fig. S2B-C). Thus, putative pUPF1-binding sites were significantly overrepresented within UPF1 targets.

**Figure 2.**
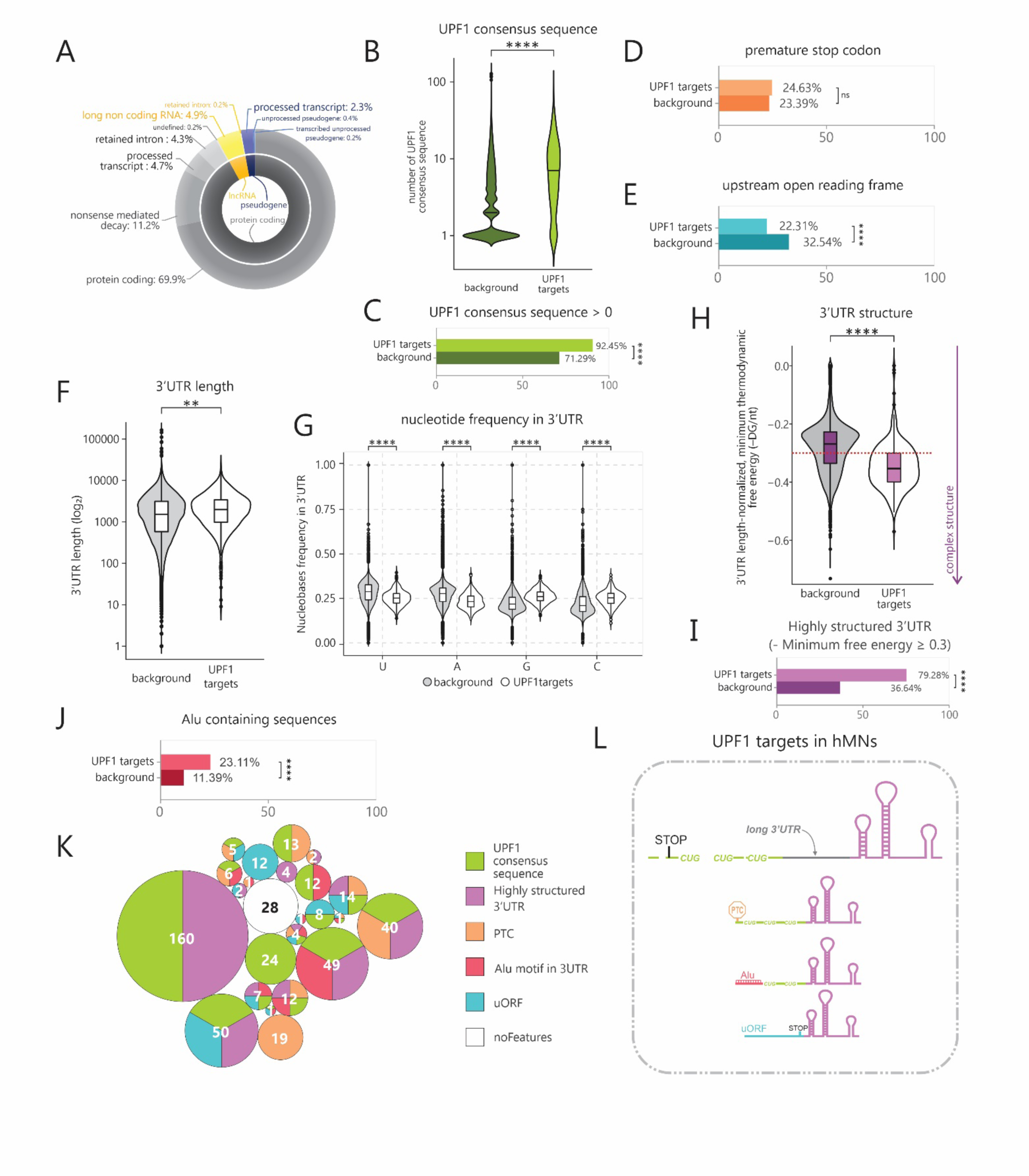
UPF1 Target Genes in MNs Exhibit Long and Highly Structured 3’UTRs. **A)** Donut chart showing the ENSEMBL annotation of the genes (inner circle) and of the transcripts (outer circle) identified as UPF1-targets in hMNs. **B)** Violin plot showing the number of consensus sequences in the 3’UTR of transcripts identified as UPF1-targets in hMNs and non-UPF1 targets (background) (t.test, p<0.0001). **C)** Bar plot showing the percentage of UPF1-target transcripts and non-UPF1 targets displaying at least one UPF1 binding sequence in the 3’UTR (chi-squared test, p<0.0001). **D)** Bar plot showing the percentage of UPF1-target transcripts and non-UPF1 targets harboring a premature stop codon (chi-squared test, not significant). **E)** Bar plot showing the percentage of UPF1-target transcripts and non-UPF1 targets presenting upstream open reading frames (chi-squared test, p<0.0001). **F)** Violin plot showing the length of 3’UTR of transcripts identified as UPF1-targets in hMNs and non-UPF1 targets (t.test, p<0.01). **G)** Violin plots showing the frequency of each nucleotide in the 3’UTR of transcripts identified as UPF1-targets in hMNs and non-UPF1 targets (t.test, p<0.0001). **H)** Violin plot showing the 3’UTR length-normalized minimum thermodynamic free energy of transcripts identified as UPF1-targets in hMNs and non-UPF1 targets. Red line=-0.3 (t.test, p<0.01). **I)** Bar plot showing the percentage of UPF1-target transcripts and non-UPF1 targets with highly structure 3’UTR (minimum free energy≥.03) (chi-squared test, p<0.0001). **J)** Bar plot showing the percentage of UPF1-target transcripts and non-UPF1 targets presenting an Alu motif in the 3’UTR (chi-squared test, p<0.0001). **K)** Circular treemap displaying the overlapping features of UPF1-target transcripts and non-UPF1 targets in hMNs. **L)** Schematic representation of the most prevalent features observed in UPF1-target transcripts in hMNs.

The most well-described UPF1-dependent RNA degradation pathway of NMD involves mRNAs retaining an exon-junction complex (EJC) downstream of a translation termination codon, whether it be a normal termination codon or a PTC (Buchwald et al., 2010; Hosoda et al., 2005). Surprisingly, we found that only about 25% of the UPF1-targets in MNs harbored a PTC, a percentage that did not significantly differ from the background (Fig. 2D and Fig. S2D). Additionally, only about 22% encoded an upstream open reading frame (uORF), a potential trigger for EJC-dependent NMD, and in fact transcripts encoding an uORF were significantly under-represented within UPF1 targets (Fig. 2E and Fig. S2E).

UPF1-associated RNA decay can also target transcripts without a 3ʹ UTR EJC, in a mechanism described as 3ʹUTR EJC-independent NMD, as a long 3ʹ UTR can increase the probability of UPF1 binding downstream of the normal termination codon and undergoing phosphorylation and activation (Hurt et al., 2013; Kurosaki et al., 2019; Singh et al., 2008). Importantly, we found that the length of the 3’UTRs was significantly longer (Fig. 2F), as well as the frequency of G and C nucleobases within 3’UTRs was significantly higher, within UPF1 targets (Fig. 2G). Sequence-specific features within the 3’UTR of mRNAs such as base pairing can lead to structure-mediated RNA degradation (SRD) through a mechanism requiring both UPF1 and G3BP1 (Fischer et al., 2020). To directly investigate 3’UTR structure, we calculated the length-normalized, minimum thermodynamic free energy (–ΔG/nt) (Zubradt et al., 2017) and found that the vast majority of UPF1-target transcripts we identified in MNs had a highly structured 3’UTR with a minimum free energy equal to or higher than 0.3 (79% v. 37%) (Fig. 2H-I). Interestingly, PTC-containing transcripts also showed a highly structured 3’UTR in addition to the presence of PTC (Fig. S2F).

Staufen-mediated decay (SMD), another UPF1 degradation pathway that depends on the double-stranded RNA-binding protein Staufen is thought to be triggered by RNA primary sequence, specifically Alu elements within the 3’UTR, which can base-pair in trans with another Alu element to form double stranded RNA structures (Park and Maquat, 2013). Our analysis showed that while only 23% of the transcripts targeted by pUPF1 contained known Alu repeat sequences within their 3’UTR, this was a substantially larger percentage relative to the background, suggesting that SMD could be a significant contributing degradation pathway utilized by MNs (Fig. 2J). Lastly, we found no differences in the length of the ORF (Fig. S2G), while 5’UTRs appeared to be shorter and contained a higher frequency of C and G nucleotides (Fig. S2H-I).

Many of the annotated UPF1 targets exhibit overlapping features (Fig. 2K-L). More than half of the PTC-or uORF-containing transcripts also include a highly structured 3’UTR. Some transcripts appear to have the potential to undergo different RNA degradation pathways, such as NMD, SMD or SRD, while only 10% lack all these structural characteristics (Fig. 2K-L). Collectively, our data demonstrate that in human MNs the length and the folding structure of the 3’UTR are the predominant characteristics of pUFP1-target mRNAs, that are degraded through various decay pathways.

### UPF1-Dependent RNA Degradation Enhances Autophagy in Human MNs

We next focused on characterizing the functional features of the proteins encoded by the transcripts targeted by pUPF1 in human MNs. Gene ontology and functional interconnection analyses revealed a strong enrichment of genes belonging to specific categories, including “lysosomal” and “ER compartments” for cellular component, “helicase activity” and “ubiquitin-related annotations” for molecular function, as well as “nucleocytoplasmic transport”, “RNA splicing” and “autophagy” (Fig. 3A and Fig. S3A-B). Autophagy was the strongest KEGG-enriched pathway, encompassing 16 genes including 13 protein-coding isoforms and 3 transcripts harboring PTCs that do not result in functional proteins (Fig. 3B). We traced back the post-transcriptional regulation of these mRNAs by pUPF1 in our RNA-seq analysis (Fig. 3C and Fig. S3C) and performed qRT-PCR with specific primers to confirm the increased mature mRNA to pre-mRNA ratio upon UPF1 KD in independent differentiations, validating successfully 9/10 targets tested (Fig. 3D and Fig. S3D). Furthermore, we selected 3 autophagy genes and tested whether the increase in their mRNA level following inhibition of UPF1-mediated RNA degradation would lead to higher protein levels. We knocked down UPF1 in three independent MN differentiations, collected samples after 40 days and performed western blot (WB) analysis. Both RPTOR and ATG4B were upregulated upon UPF1 KD, while ATG12, for which the transcript targeted by UPF1 harbors a PTC, was not affected (Fig. 3E). The strong enrichment for genes involved in this pathway suggests that MNs may utilize UPF1-dependent RNA decay as a mechanism to modulate autophagic flux.

**Figure 3.**
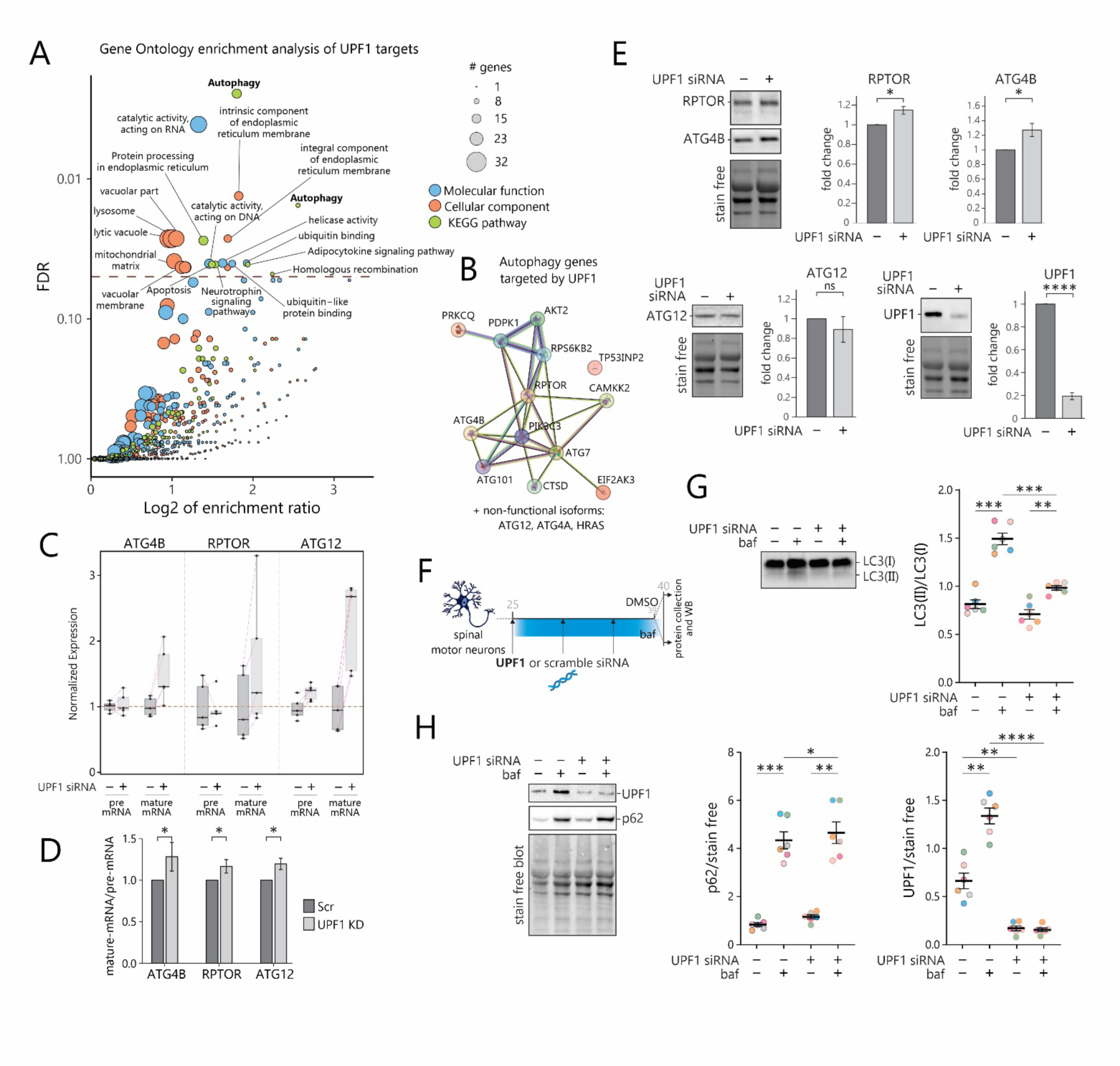
UPF1-Dependent RNA Degradation Modulates Autophagy in Human MNs. **A)** GO Enrichment analysis of the UPF1 targets genes. The dashed line indicates the FDR value of 0.05. The size of the circles is proportional to the number of genes of indicated categories. In blue are highlighted the groups associated with the Molecular Function dataset, in orange those with the Cellular Component dataset and in green those with the KEGG pathway. **B)** Functional and physical association graph of proteins involved in autophagic mechanisms identified as UPF1 targets. Line color indicates the type of interaction evidence. (Known interactions: Light blue, from curated databases; Purple, experimentally determined. Predicted Interactions: Green, gene neighborhood. Red, gene fusions. Blue, gene co-occurrence). **C)** Boxplots showing pre-mRNA and mature mRNA modulation of autophagy genes *ATG4B*, *RPTOR* and *ATG12* gene after treatment with siRNA targeting *UPF1* from RNA-seq data. Pink dotted lines connect samples from the same differentiation (n=5 independent differentiations). **D)** Bar plot showing the fold change of the pre-mRNA/mature mRNA ratio of *ATG4B*, *RPTOR* and *ATG12* in MNs treated with scramble or UPF1 siRNA, analyzed by qPCR (3 to 6 independent differentiations, paired t.test, *p<0.05). **E)** Representative WB bands and bar plots showing the fold change of RPTOR/raptor, ATG4B, ATG12 and UPF1 proteins in hMNs treated with scramble or UPF1 siRNA. Protein levels are normalized to protein loading (n=3 independent differentiations; t.test, *p<0.05; ****p<0.0001, ns=not significant). **F)** Experimental schematic used to assess autophagy flux by treating hMNs with DMSO or bafilomycin in UPF1 KD or control conditions. **G)** Left: Representative WB bands for LC3(I) and LC3(II) in hMNs treated with DMSO or bafilomycin in UPF1 KD or control conditions. Right: bar plot showing LC3(II)/LC(I) ratio in hMNs treated with DMSO or bafilomycin in UPF1 KD or control conditions (n=6 biological replicates from 2 independent differentiations; paired t.test, **p<0.01; ***p<0.001). **H)** Left: Representative WB bands for UPF1 and p62 in hMNs treated with DMSO or bafilomycin in UPF1 KD or control conditions. Right: bar plot showing UPF1 and p62 levels in hMNs treated with DMSO or bafilomycin in UPF1 KD or control conditions. Protein levels are normalized to protein loading (n=6 biological replicates from 2 independent differentiations; paired t.test, **p<0.01; ***p<0.001; ****p<0.0001).

Critically, UPF1 targets regulatory proteins that enhance the autophagy pathway (e.g., ATG4B: autophagy related 4B cysteine peptidase) (Maruyama and Noda, 2017; Yamamoto et al., 2023), or inhibit it (e.g., RPTOR: regulatory-associated protein of mTOR) (Khalil et al., 2023). To determine whether the autophagic flux is regulated positively or negatively by UPF1, we treated iPSC-derived MNs for 24 hours with the autophagy blocker bafilomycin in cultures pre-treated with *UPF1* siRNA or siRNA-scrambled control (Fig. 3F). We then collected protein and quantified the levels of the autophagy markers LC3 and p62 by WB. As expected, treatment with bafilomycin induced an increase in the LC3(II)/LC3(I) ratio, as well as an accumulation of p62 (n=6, p<0.01) in both conditions (Fig. 3G-H). At the same time, MNs depleted of UPF1 exhibited a substantial reduction in LC3(II) accumulation, suggesting defects in autophagosome formation or autophagosome-lysosome fusion (n=6, p=0.007) (Fig. 3G). We also found a moderate but significant increase in p62 levels in UPF1-KD MNs relative to controls before and after bafilomycin treatment, reinforcing the hypothesis that UPF1 increases the efficiency of the autophagic flux in healthy MNs (n=6, p=0.043) (Fig. 3H). Intriguingly, blocking autophagy caused a dramatic upregulation of UPF1 protein in control conditions, further suggesting an interplay between the autophagy and UPF1-dependent RNA decay pathways (Fig. 3H).

### TDP-43 Loss of Function Causes a Reduction in UPF1-Dependent mRNA Degradation

ALS MNs exhibit compromised RNA metabolism, particularly associated with TDP-43 dysfunction (Archbold et al., 2018; Barmada et al., 2010; Donde et al., 2019; Klim et al., 2019). However, the activity of UPF1-dependent mRNA degradation pathways in the context of ALS remains controversial (Barmada et al., 2015; Held et al., 2023; Jackson et al., 2015; Kamelgarn et al., 2018; Ortega et al., 2020; Sun et al., 2020; Xu et al., 2019; Zaepfel et al., 2021). Using the *bona fide* list of mRNAs targeted by pUPF1 in MNs, we next sought out to determine whether and how RNA decay pathways are affected in ALS. We first used a dataset generated from laser-captured lower MNs of postmortem *C9ORF72* ALS patient and healthy control tissue (Cooper-Knock et al., 2015) and performed gene set enrichment analysis (GSEA) to assess the expression of UPF1 targets as a class. This analysis showed that transcripts modulated post-transcriptionally by pUPF1 were significanlty upregulated as a group (adj. p=0.03) (Fig. 4A), suggesting a reduction in either the efficiency of degradation, or the activity of UPF1 pathways. This stands in agreement with a recent analysis in MNs dissected from sporadic ALS patients (Krach et al., 2018,Held, 2023 #77).

**Figure 4.**
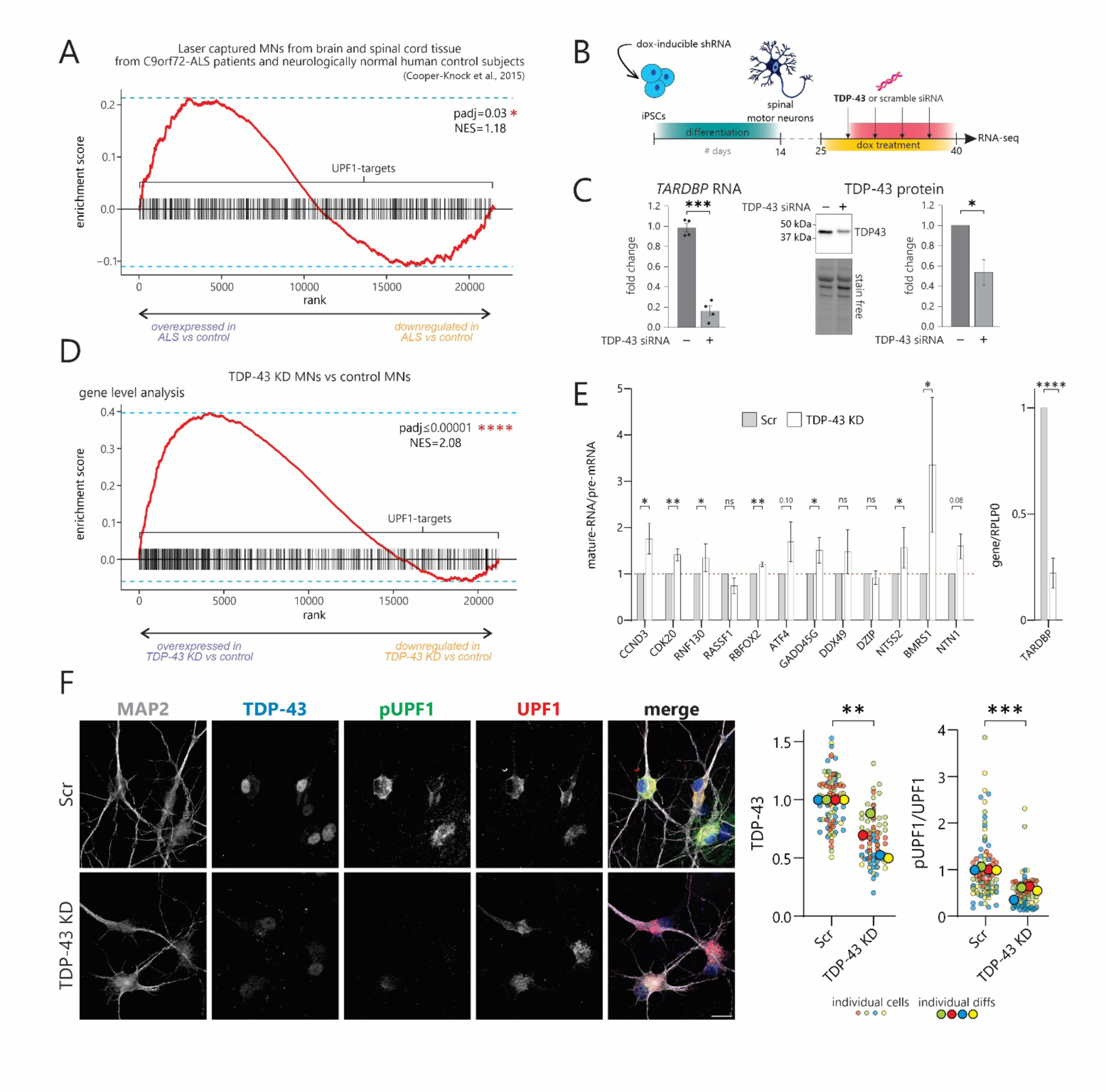
TDP-43 Loss of Function Causes a Reduction in UPF1 Dependent mRNA Degradation. **A)** Enrichment plot from GSEA of laser captured MNs from brain and spinal cord tissue from *C9ORF72* ALS patients and neurologically normal individuals(Cooper-Knock et al., 2015). The plot contains the profile of the running enrichment score (ES) and positions of gene set elements on the rank ordered list in GSEA. The gene set includes hMNs UPF1-target genes. (NES=Normalized Enrichment Score). **B)** Experimental schematic used to knock down TDP-43 in iPSC-derived hMNs. **C)** Left: Bar plot showing the RNA level of *TARDBP* in hMNs after TDP-43 KD treatment, analyzed by qPCR (p<0.001). Right: representative WB for TDP-43 in MN cultures after TDP-43 KD treatment. Right: bar plot showing the fold change in the TDP-43/loading control ratio after TDP-43 KD treatment (t.test, p<0.05). **D)** Enrichment plot from GSEA of TDP-43 KD and scramble siRNA-treated hMNs. The plot contains the profile of the running enrichment score (ES) and positions of gene set elements on the rank ordered list in GSEA. The gene set includes hMNs UPF1-target genes. (NES=Normalized Enrichment Score). **E)** Left: Bar plot showing the fold change of the pre-mRNA/mature mRNA ratio of selected UPF1-target genes in TDP-43 KD and scramble siRNA-treated hMNs treated, analyzed by qPCR. Right: Bar plot showing the fold change of *TARDBP* after treatment with doxycycline and siRNA, analyzed by qPCR and normalized to the level of housekeeping gene RPLP0 (3 to 6 independent differentiations, paired t.test, *p<0.05; **p<0.01; ****p<0.0001). **F)** Left: representative confocal microscopy images of TDP-43 KD and scramble siRNA-treated hMNs immunolabeled for TDP-43 (blue), pUPF1 (green), total UPF1 (red) and MAP2 (white). Dashed lines mark the neuronal soma. Scale bars represent 25 µm. Right: SuperPlot displaying the fold change of TDP-43 and pUPF1/total UPF1 signal in TDP-43 KD and scramble siRNA-treated hMNs. Small dots represent individual cells. Big dots represent individual differentiations and were used to run the statistical test. Colors highlight each individual differentiation (paired t.test, *p<0.01; ***p<0.001).

To more directly investigate the impact of TDP-43 dysfunction on UPF1-dependent RNA decay, we next established an iPSC-based MN model where we could effectively modulate TDP-43 levels. We generated a stable iPSC line containing a doxycycline-inducible shRNA targeting TDP-43 and then induced its differentiation to spinal MNs as described above (Ziller et al., 2018). By day 40, the cultures exhibited a substantial reduction of TDP-43 at both RNA (80%, p<0.001) and protein levels (50%, p<0.5) (Fig. 4B-C). We then performed RNA-seq across four independent samples from two differentiations. GSEA showed a very substantial effect where both pUPF1 target-genes (Fig. 4D) and transcripts (Fig. S4A) were significantly upregulated as a group after TDP-43 loss-of-function (adj. p <0.00001). We verified that the higher expression level was driven by UPF1-dependent post-transcriptional regulation using qRT-PCR for 8/12 genes that we tested (Fig. 4E). Lastly, by performing immunocytochemistry for UPF1 and pUPF1 in these MN cultures, we found that KD of TDP-43 caused a substantial reduction in UPF1 phosphorylation, as measured by the pUPF1/UPF1 ratio (n=4, p value <0.001) (Fig. 4F). Collectively, these results demonstrate that TDP-43 loss-of-function in MNs causes a reduction in UPF1-dependent mRNA decay activity.

### The Degradation of TDP-43 Dependent PTC-Containing Transcripts Requires UPF1 and UPF2

The disruption of TDP-43 activity in MNs results in gross mis-splicing of mRNAs (Brown et al., 2022; Klim et al., 2019; Ma et al., 2022; Tank et al., 2018) and progressive buildup of novel species with cryptic or skiptic exons, some of which encode PTCs (Brown et al., 2022; Ma et al., 2022). We next sought to determine whether these TDP-43 dependent mis-splicing events constitute novel substrates for UPF1 degradation. We first systematically characterized the nature of mRNA metabolism defects in our spinal MN cultures depleted of TDP-43. Applying differential splicing analysis tools (Vaquero-Garcia et al., 2016) revealed 745 TDP-43 dependent, significantly mis-spliced events associated with 528 genes, including previously reported *STMN2*, *UNC13A*, *UNC13B* and *KCNQ2* transcripts (Fig. 5A). The majority of these mis-spliced RNAs were significantly downregulated in their expression (GSEA, adj. p <0.00001) (Fig. 5B-C), suggesting that they were subject to degradation mechanisms. This group includes mis-spliced mRNAs with PTCs that would likely be subject to UPF1-dependent NMD that exhibit a clear and significant downregulation in expression (31/81 genes are significantly downregulated, GSEA, adj. p <0.00001) (Fig. 5E and S5A-B). To directly assess the role of UPF1 on the degradation of these PTC-containing mRNAs, we simultaneously knocked down TDP-43 and UPF1, or TDP-43 alone, in MN cultures over the course of 15 days and performed another round of RNA-seq (Fig. 5D and S5C). Surprisingly, the expression of TDP-43 dependent PTC-containing mRNAs was not strongly affected by the downregulation of UPF1 and consequent inhibition of NMD (GSEA for TDP-43 KD *vs*. TDP-43 and UPF1 KD, adj. p=0.22) (Fig. 5F and S4D-E). Consistent with a lack of an effect, applying MAJIQ-based splicing analysis within the double TDP-43/UPF1 KD conditions did not reveal a preferential increase in PTC-containing cryptic transcripts relative to TDP-43 KD alone, except for 6 transcripts (Fig. 5G and Fig. S5F). As an example, there was no increase in cryptic PTC-containing *UNC13A* mRNA (Fig. 5H, top), or *STMN2* mRNA, which does not harbor a PTC (Fig. 5H, bottom), whereas *PFKP* represented one the few (6/86) PTC-containing transcripts that increased upon double TDP-43/UPF1 KD (Fig. S5F).

**Figure 5.**
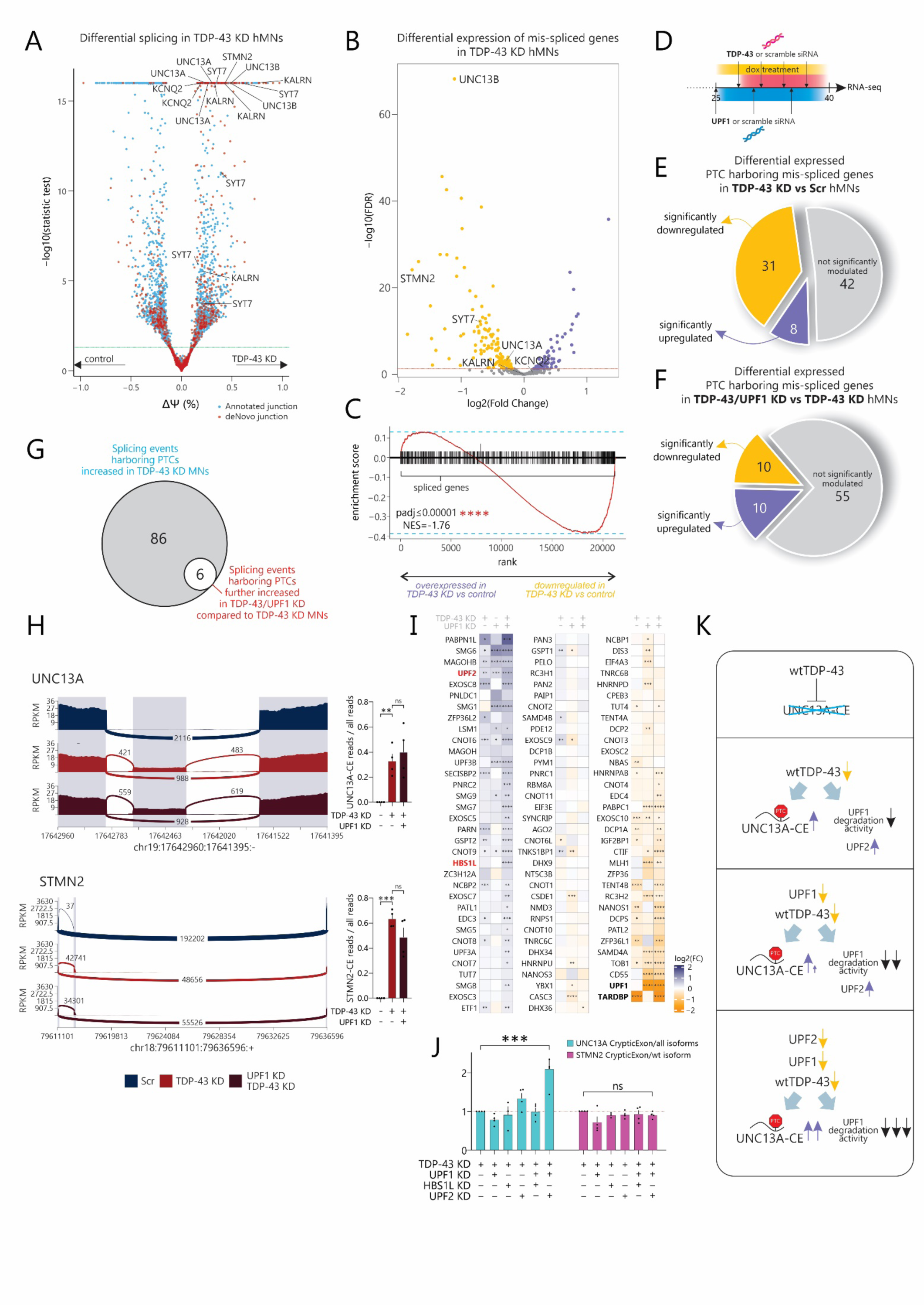
The Degradation of TDP-43 Dependent PTC-Containing Transcripts Requires UPF1 and UPF2. **A)** Volcano plot showing differential splicing analysis by MAJIQ(Vaquero-Garcia et al., 2016) in TDP-43-KD and scramble siRNA-treated iPSC-derived hMNs (n=4 biological replicates from 2 individual differentiations). Each point represents a splice junction. **B)** Volcano plot showing differential expression analysis of TDP-43 splicing targets in TDP-43-KD and scramble siRNA-treated iPSC-derived hMNs (n=4 biological replicates from 2 individual differentiations). **C)** Enrichment plot from GSEA of TDP-43 KD and scramble siRNA-treated hMNs. The plot contains the profile of the running enrichment score (ES) and positions of gene set elements on the rank ordered list in GSEA. The gene set includes TDP-43 splicing targets genes. (NES=Normalized Enrichment Score). **D)** Experimental schematic used to knock down both TDP-43 and UPF1 in iPSC-derived hMNs. **E)** Pie chart displaying the proportion of TDP-43 dependent mis-spliced genes harboring PTCs that are significantly up-or down-regulated in hMNs after TDP-43 KD. **F)** Pie chart displaying the proportion of TDP-43 dependent mis-spliced genes harboring PTCs that are significantly up-or down-regulated in TDP-43/UPF1 KD condition compared to TDP-43 KD condition. **G)** Euler plot showing the proportion of TDP-43 dependent splicing events harboring PTCs that increase in TDP-43/UPF1 KD condition compared to TDP-43 KD condition. **H)** Left: Sashimi plots showing mean cryptic event occurrence in *UNC13A* and *STMN2* in scramble siRNA treated, TDP-43 KD and TDP-43/UPF1 KD hMNs. Right: Bar plot showing quantification of CE-containing reads/total reads ratio of *UNC13A* and *STMN2* from RNA-seq of scramble siRNA treated, TDP-43 KD and TDP-43/UPF1 KD conditions (n=4 biological replicates from 2 individual differentiations; t.test, **p<0.01; ***p<0.001, ns=not significant). **I)** Heatmap showing the average expression level of genes implicated in RNA-decay pathway mechanism, plus TARDBP, upon UPF1, TDP-43 and TDP-43/UPF1 KD conditions over scramble-siRNA treated hMNs by RNA-seq (*FDR< 0.05, **FDR<0.01, ***FDR<0.001, **** FDR<0.0001). **J)** Bar plot showing the fold change of UNC13A-CE/total UNC13A (light blue) and STMN2-CE/wt STMN2 (fuchsia) analyzed by qPCR in TDP-43-KD hMNs treated with siRNA targeting UPF1, HBS1L and UPF2 as indicated in the figure (n=4 biological replicates from 2 individual differentiations, One-way ANOVA followed by Dunnett multiple comparison test, ***p<0.001, ns=not significant). **K)** Schematic representation of our working model. TDP-43 loss of function leads to mis-splicing and accumulation of PTC-containing transcripts, and to downregulation of UPF1-dependent mRNA degradation activity. In TDP-43/UPF1 KD there is a mild effect on the production of CE-containing transcripts, that become conspicuous in a TDP-43/UPF1/UPF2 condition.

We thus next asked whether TDP-43-dependent PTC-containing mRNAs are degraded by other pathways or if compensatory mechanisms come into play once NMD is impaired. Examining the expression of genes encoding proteins involved in RNA decay mechanisms revealed a significant upregulation of several candidates including *UPF2* and *HBS1L* (Fig. 5I and Fig. S5G). HBS1L is a protein involved in non-stop decay (NSD) and no-go decay (NGD), pathways designated for the identification and degradation of mRNAs that have stalled ribosomes during translation elongation (Doma and Parker, 2006; Frischmeyer et al., 2002), while UPF2 is a key player in NMD and contributes to the stability and assembly of the UPF1 complex on the mRNA designated for degradation on subsets of degradation targets (Huang et al., 2011; Yepiskoposyan et al., 2011). Close examination of the expression of these two genes showed that while both *HBS1L and UPF2* were upregulated when TDP-43 and UPF1 were simultaneously knocked down, *UPF2* levels were also elevated when TDP-43 alone was depleted (Fig. 5I and Fig. S5G). To test the potential contribution of these factors to mRNA degradation we differentiated MNs, treated them with an antisense oligonucleotide (ASO) targeting *TDP-43,* and at the same time used an siRNA to knock down *HBS1L* or *UPF2* either alone, or together with *UPF1*. We confirmed the efficient reduction in the expression of these factors in each condition by qRT-PCR (Fig. S5H-I) and quantified the abundance of specific TDP-43-dependent mRNAs, i.e. *UNC13A* and *STMN2*. The relative expression of cryptic PTC-containing *UNC13A* was exclusively increased when both *UPF1* and *UPF2* were knocked down, while reduction in each one of the three factors alone had no effect (Fig. 5J), while *STMN2* was not affected by any of these additional genetic manipulations (Fig. 5J). These data suggest that the degradation of some of the TDP-43-dependent PTC-containing mRNAs requires the combinatorial actions of UPF1 and UPF2 (Fig. 5K).

### UPF1 and TDP-43 Regulate the Polyadenylation and 3’UTR Length of Select Target Transcripts

Given our finding that a dominant trait of UPF1-targets in MNs is a long 3’UTR, we next sought to investigate whether TDP-43 loss-of-function disrupts alternative polyadenylation (APA) and consequently the 3’UTR length. TDP-43 has been reported to modulate the length of 3’UTRs by adjusting the selection of APA sites in specific genes such as *SOX2* and *STMN2,* as well as within *TDP-43* mRNA itself as a means of autoregulation (Ayala et al., 2011; Melamed et al., 2019; Modic et al., 2019). However, the broader role of this mechanism on MN transcripts has not been demonstrated. Application of <u>D</u>ynamic <u>a</u>nalyses of Alternative <u>P</u>oly<u>A</u>denylation from <u>R</u>NA-<u>s</u>eq (DaPars) (Xia et al., 2014) in our 40-day old spinal MN cultures depleted of TDP-43 revealed 2,033 recurrent alternatively polyadenylated target transcripts relative to controls (Fig. 6A), demonstrating a very significant effect on APA site selection. Approximately 73% of these events resulted in longer 3′UTRs, while 27% caused shortening (Fig. 6A). Importantly, 34% of these events occur in regions showing binding sites for TDP-43 in iCLIP data generated in HEK293 cells (Fig. 6B), suggesting a direct effect (Rot et al., 2017). The length of the 3’UTR can influence the stability and expression of RNA (Agarwal et al., 2015; Lianoglou et al., 2013; Tushev et al., 2018), through several mechanisms that are partially cell type-specific (Chen et al., 2018; Tushev et al., 2018). Transcripts with longer 3’UTRs appeared to be moderately but significantly downregulated as a group (GSEA, adj. p=0.00004) (Fig. 6C), while no changes in expression were observed for transcripts with shorter 3’UTRs (GSEA, adj. p=0.14) (Fig. S6A). Beyond stability, APA can also affect the rate of translation (Mayr, 2016), as well as the trafficking and localization of mRNAs (Berkovits and Mayr, 2015; Horste et al., 2023). To assess the functional enrichment of TDP-43 targets that are regulated by APA, we performed GO analysis on transcripts with a longer 3’UTR in TDP-43-depleted MNs. We found these to be very strongly enriched for genes regulating “synaptic vesicle transport and localization” and “regulation of neuromorphogenesis” (Fig. S6B), as well as for proteins residing in axons, presynaptic and synaptic vesicles, and microtubules (Fig. 6D). These data suggest that TDP-43 depletion results in disruptions in APA site selection and causes the accumulation of mRNAs with longer 3’UTRs that encode proteins with key functional roles in neuronal physiology.

**Figure 6.**
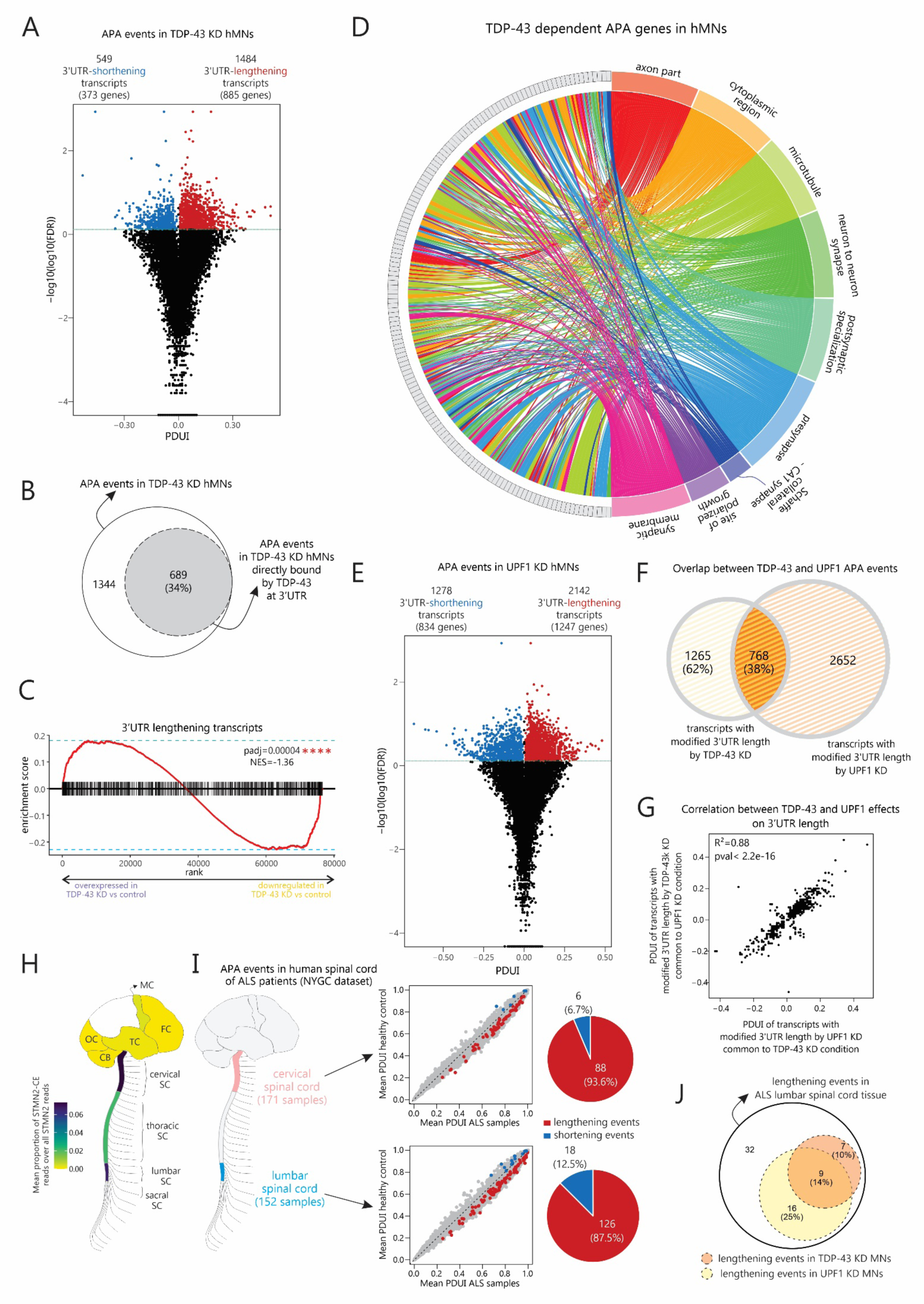
UPF1 and TDP-43 Regulate the Polyadenylation and 3’UTR Length of Select Target Transcripts. **A)** Iceberg plot showing APA events in TDP-43 KD hMNs. **B)** Euler plot showing APA events in TDP-43 KD hMNs that have been shown to be directly bound by TDP-43 at 3’UTR in iCLIP experiment (Rot et al., 2017) **C)** Enrichment plot from GSEA of TDP-43 KD and scramble siRNA-treated hMNs. The plot contains the profile of the running enrichment score (ES) and positions of gene set elements on the rank ordered list in GSEA. The gene set includes hMNs genes displaying 3’UTR lengthening events upon TDP-43 KD. (NES=Normalized Enrichment Score). **D)** Chord plot displaying the significant enrichments of GO analysis (Cellular Component not redundant dataset) associated with APA genes resulting in longer 3’UTR in TDP-43 KD hMNs. **E)** Iceberg plot showing APA events in UPF1 KD hMNs. **F)** Venn diagram showing the intersection of APA events upon TDP-43 KD and UPF1 KD hMNs. Percentages indicate the proportion of TDP-43 events overlapping or not overlapping UPF1 events. **G)** Dot plot showing the correlation between TDP-43 KD and UPF1 KD effect on 3’UTR length of events occurring upon both conditions. **H)** Schematic illustrating TDP-43 dysfunction level different central nervous system regions from ALS patients calculated as the mean proportion of *STMN2*-CE reads/total *STMN2* reads from NYGC repository (MC=Motor Cortex; FC=Frontal Cortex; OC=Occipital Cortex; TC=Temporal Cortex, CB=Cerebellum; SC=Spinal Cord). **I)** Left: schematic illustration of the CNS regions analyzed from NYGC repository. Right: Dot plots showing the APA events occurring in human cervical (top) and lumbar (bottom) spinal cord of ALS patients and pie charts showing the proportion of significant lengthening (red) and shortening (blue) events. **J)** Euler plot showing the APA events occurring in lumbar spinal cord of ALS patients that we observed by knocking down TDP-43 (yellow circle) or UPF1 (orange circle) in iPSC-derived hMNs.

Intriguingly, our previous genomics analyses of pUPF1-mediated regulation in MNs highlighted the length of the 3’UTR as a predominant trait of target transcripts (Fig. 2). To determine whether UPF1 loss-of-function would also cause the accumulation of alternatively polyadenylated transcripts we applied DaPars on 40-day old spinal MN cultures depleted of UPF1. This analysis identified 3,420 transcripts with APA sites, with 63% of them exhibiting 3′ UTR lengthening and 37% shortening, demonstrating a substantial effect of UPF1 KD (Fig. 6E). Critically, at least 45% of these targets appeared to be bound to pUPF1 in the previous RIP-seq experiment (Fig. S6C). Suprisingly, when we compared the two datasets we found that TDP-43 and UPF1 depletion impacts the APA and 3′UTR length of the same 768 transcripts, representing a substantial 38% of all TDP-43 targeted transcripts and 45% of all TDP-43 targeted genes (Fig. 6F and Fig S6D). This overalp was deemded highly significant by hypergeometric statistical testing (p<1e^-20^). Moreover, the overlapping transcripts appeared to be similarly affected by TDP-43 and UPF1 KD, exhibiting concordant longer or shorter 3’UTRs in each one of the two independent KD conditions (Fig. 6G). Collectively, these data demonstrate the unexpected finding that UPF1 and TDP-43 exhibit overlapping effects on alternative polyadenylation and regulation of 3’UTR length of select target transcripts in MNs that are strongly enriched for proteins with axonal and synaptic functions (Fig. S6E and S6F).

To investigate whether the APA changes we identified in iPSC-derived MNs *in vitro* reflect changes that occur in ALS patients with TDP-43 pathology we performed DaPars on a large publicly available NYGC dataset (207 ALS patient samples) of RNA-seq from postmortem CNS tissue subtypes including brain and spinal cord sections. As a surrogate of TDP-43 dysfunction we first measured the abundance of mis-spliced *STMN2*, as an established molecular marker of TDP-43 neuropathology (Fig. 6H). This analysis showed that the cervical and lumbar spinal cord were the most affected tissues in the ALS patient population including C9ORF72 and sporadic individuals. DaPars on the cervical and lumbar spinal cord bulk RNA-seq samples of ALS patients revealed respectively 94 and 144 transcripts that undergo significantly different APA relative to healthy controls (Fig. 6I), with the large majority of these events resulting in longer 3′ UTRs (Fig. 6I). Among the 64 genes showing 3’UTR lengthening events in lumbar spinal cord, 16 were also found in TDP-43 depleted iPSC-derived MNs, 25 were also found in UPF1 depleted iPSC-derived MNs, and 9 were overlapping in both datasets, suggesting co-regulation (Fig. 6J). We further explored the NYGC dataset for APA events, performing separate analysis on samples harboring *C9ORF72* mutation and samples from sporadic ALS patients (Fig. S6G). Interestingly, there was a large concordance between the two groups (Fig. S6H), with more APA events found in sporadic samples (Fig. S6G and Fig. S6H), although this could reflect different sample sizes. Finally, we used RNA-seq data from TDP-43 positive, and TDP-43 negative neuronal nuclei isolated from frontal cortex tissue of C9ORF72 ALS/FTLD patients(Liu et al., 2019) to assess APA changes. We observed 41 events resulting in longer 3’UTR and 63 events causing 3’UTR shortening (Fig. S6I). Among the genes showing longer 3’UTR, 6 showed the same effect in TDP-43 depleted MNs and 6 in UPF1 depleted MNs *in vitro* (Fig. S6J). Three genes were overlapping in both datasets, and two of them share the same 3’UTR lengthening phenotype also in NYGC samples. Collectively, these results suggest that TDP-43 modulates APA site selection and UPF1 selectively degrades a substantial proportion of TDP-43-dependent transcripts that end up with longer 3’UTRs as a result of ALS-associated TDP-43 dysfunction.

## DICUSSION

The interplay between UPF1-dependent mRNA decay pathways and ALS pathophysiology is complex. The dysfunction of TDP-43, which characterizes the majority of ALS patients (Chen-Plotkin et al., 2010; Neumann et al., 2006), results in a disruption in the normal splicing of several target mRNAs some of which encode PTCs (Akiyama et al., 2022; Klim et al., 2021; Mehta et al., 2023). Whether or not these constitute UPF1 substrates and how their accumulation impacts mRNA surveillance mechanisms in MNs has remained unknown. At the same time, overexpression of UPF1 has been shown to mitigate ALS neurotoxicity associated with mutant TDP-43 and C9ORF72 *in vitro* and *in vivo*; however the precise mechanisms underlying this effect has been a topic of debate, despite the fact that UPF1 has been developed as a potential gene therapy for ALS (Barmada et al., 2015; Jackson et al., 2015; Ortega et al., 2020; Xu et al., 2019; Zaepfel et al., 2021). Here we sought out to address these outstanding questions by first defining a highly stringent set of mRNAs that UPF1 modulates under physiological conditions in MNs, which are the most susceptible cells to ALS disease mechanisms.

The state of NMD has been previously investigated in different cellular models of *C9ORF72* and ALS patient tissue by examining the expression of a few UPF1 target genes with conflicting results (Held et al., 2023; Xu et al., 2019; Zaepfel et al., 2021). Our approach shifted from focusing on individual genes, which may be susceptible to various levels of cell type-specific transcriptional and posttranscriptional regulation, to analyzing the expression of a large group of high-confidence, and MN-specific UPF1 target genes. We identified these by intersecting datasets from RIP-seq for phosphorylated UPF1 (Kurosaki et al., 2014), with post-transcriptional gene expression changes detected in MNs depleted of UPF1. This stringent set of criteria ensured that we identified direct targets of pUPF1 activity, which is particularly important given that UPF1 is known to promiscuously bind the genome and become activated by SMG1 phosphorylation only when a gene is subject to degradation (Kurosaki et al., 2019). We surprisingly found that UPF1 target mRNAs in MNs were not enriched for PTC-containing transcripts, but rather for transcripts with especially long 3’UTRs. Using the expression of these *bona fide* targets as a biomarker of UPF1-dependent activity we found a clear and strong negative effect of TDP-43 loss-of-function on the efficiency of UPF1-dependent mRNA surveillance. This was further supported by a significant downregulation in the level of UPF1 phosphorylation in concomitant MN cultures, as well as in a dataset of ALS-C9ORF72 patient MNs isolated by laser capture of spinal cord tissue (Cooper-Knock et al., 2015). Our conclusions are in line with a recently reported dysregulation of NMD in another laser-captured MN dataset from sporadic ALS patients (Held et al., 2023), as well as in L5 extratelencephalic neurons identified through single cell RNA-seq in postmortem ALS patient tissue (Gittings et al., 2023).

The compromised UPF1 activity offers a plausible rationale for the considerable accumulation of transcripts containing PTCs in neurons with dysfunctional TDP-43 (Brown et al., 2022; Ma et al., 2022), which should typically be degraded by NMD quite efficiently (Ge and Porse, 2014; Lewis et al., 2003). Intriguingly, we found that further impairing NMD by knocking down *UPF1* in addition to *TDP-43* resulted in a pronounced upregulation of only a handful of PTC-containing transcripts, with most unaffected. However, the downregulation of UPF2, a core component of the NMD machinery, which binds to UPF1 and enhances its catalytic activity on mRNA substrates (Kurosaki et al., 2019), caused a substantial increase in the abundance of mis-spliced PTC-containing transcripts like *UNC13A*. Given that our siRNA approach did not result in complete ablation of UPF1 expression, the increase in UPF2 we observed in the double TDP-43/UPF1 KD MNs may act in a compensatory manner to maintain critical mRNA surveillance levels.

The precise mechanism by which TDP-43 loss-of-function impairs UPF1 activity remains unclear and warrants further investigation. The dramatic reduction in UPF1 phosphorylation suggests a breakdown in SMG1 or AKT enzymatic activity, although it may be possible that other stages of the process are inhibited such as the binding and recognition of substrate mRNAs, particularly given the reported direct interaction between TDP-43 and UPF1 (Freibaum et al., 2010), although this has not been yet confirmed in MNs. Additionally, our finding that under physiological conditions UPF1 modulates cellular pathways such as autophagy by targeting the degradation of key regulators, posits the alternative premise that the reduction in UPF1 activity may in fact reflect a cellular response to TDP-43-dependent stress. Importantly, disruptions in autophagy and proteostasis represent fundamental facets of ALS pathophysiology (Chua et al., 2022; Webster et al., 2016). Pathological inclusions characterized by LC3 and SQSTM1 that are indicative of failed or stalled autophagic clearance, have been documented in sporadic ALS patient MNs (Sasaki, 2011). The loss of TDP-43 has been reported to enhance autophagosomal biogenesis (Xia et al., 2016), and TDP-43 has been identified as a stabilizer of RPTOR a key regulator of autophagic flux and adaptor protein of MTORC1 (Xia et al., 2016), while we report here that UPF1 targets *RPTOR* mRNA for degradation. Thus, it may be that the reduction of UPF1 activity, which inhibits the synthesis of autophagosomes, is a physiological MN response to counteract TDP-43 dysfunction.

Our finding that the dominant structural feature of UPF1 target genes in MNs was a long and highly structured 3’UTR prompted us to determine whether APA, beyond PTC-triggered decay, represented another mechanism of interplay between TDP-43 and UPF1. Indeed, while previous studies had flagged the ability of TDP-43 to modulate 3’UTR length in specific genes (Ayala et al., 2011; Prudencio et al., 2015; Rot et al., 2017), we report here an extensive disruption in APA site selection and consequently 3’UTR length that is TDP-43-dependent. Most of the changes we find in MNs favor distal polyA site usage and impact genes that play crucial roles in core neuronal activity, including synaptic function, action potential firing and microtubule dynamics, all of which are disrupted in ALS pathophysiology (Taylor et al., 2016). The length of a 3’UTR can affect stability, transport, and localization (Agarwal et al., 2015; Lianoglou et al., 2013; Tushev et al., 2018), and may be particularly critical for MNs, which necessitate the transportation and translation of RNAs at locations far from the endoplasmic reticulum. Many of these events occur in transcripts that encode a TDP-43 binding site in the 3’UTR or in the last exon, suggesting that they likely reflect primary TDP-43 dependent defects. Strikingly, we uncovered that depletion of UPF1 in MNs causes a dramatic disruption in APA, affecting almost 50% of the TDP-43 target mRNAs in a similar distal direction, demonstrating overlapping regulation by these two multi-faceted mRNA binding proteins. Our observations on UPF1 decay are in accordance with a previous report that showed UPF1 preferentially degrades mRNAs retaining longer rather than shorter 3’UTR in cancer cells (Kishor et al., 2020). Notably, we detected a considerable proportion of these APA events in postmortem ALS patient spinal cord tissue validating their relevance *in vivo*. While the reason behind the translocation of TDP-43 from the nucleus remains elusive, these findings highlight APA as another major feature of TDP-43 pathophysiology and additionally demonstrate that UPF1 targets these irregular 3’UTR events to preserve mRNA integrity and quality control in iPSC-derived MNs.

Critically, our findings are in accordance with concurrent but independent observations by Gomez et al., that identified both the accumulation of transcripts with abnormally long 3’UTRs in ALS patient-derived samples and TDP-43 dysfunction iPSC-neuronal models, as well as the effects of UPF1 on targeting APA transcripts for degradation demonstrated by a UPF1 overexpression approach.

Notably, beyond ALS, nuclear depletion and cytoplasmic accumulation of TDP-43 characterizes approximately 50% of individuals diagnosed with frontotemporal dementia and Alzheimer’s disease and recent studies have shown that these patients exhibit evidence of TDP-43 dependent mis-splicing (Agra Almeida Quadros et al., 2024; Estades Ayuso et al., 2023). Thus, APA likely represents an aspect of TDP-43 pathophysiology that is relevant across neurodegenerative diseases characterized by TDP-43 proteinopathies but like cryptic splicing, reflects some degree of neuronal subtype specificity. In conclusion, our work provides a comprehensive description of UPF1-mediated mRNA decay activity in physiological and disease mimetic MNs, reveals overlapping roles between UPF1 and TDP-43 on regulating 3’UTR length, and offers novel insight into the intricate interplay between RNA metabolism and neurodegeneration in ALS.

## ACKNOWLEDGMENTS

We are grateful to the following funding sources: US National Institutes of Health (NIH), National Institute on Neurological Disorders and Stroke (NINDS) and National Institute on Aging (NIA) R01NS104219 and NINDS 1R01NS134166 (E.K.), 1R21NS131713 (E.K.), the Les Turner ALS Foundation (E.K.) and the New York Stem Cell Foundation (E.K.) and the AFM Telethon Postdoctoral Fellowship (F.A.), NIH R01GM059614 and R35GM149268 (L.E.M.) and R21GM147719 (T.K.). E.K is a Les Turner ALS Center Investigator and a New York Stem Cell Foundation – Robertson Investigator.

## METHODS

### Stem cell cultures and motor neuron differentiation

Induced Pluripotent Stem Cells (iPSCs) were fed with mTESR^TM^1 (Stam cell Technologies) for maintenance. iPSCs were differentiated into spinal motor neurons (MNs) following a standard 2D differentiation protocol (Ortega et al., 2020). iPSCs were dissociated into single cells with Accutase and plated in mTESR^TM^1 supplemented with 10μM ROCK inhibitor (Y-27632, DNSK International) at a density of 1.2million/well of a 6-well plate. After two days, cells were washed with PBS, than fed daily with N2B27 media (50% DMEM:F12 / 50% Neurobasal media, supplemented with Non-Essential Amino Acids, Glutamax, N2, B27, all from Thermo Fischer Scientific) supplemented with 10μM SB431542 (DNSK International), 100nM LDN-193189 (DNSK International), 1μM Retinoic Acid (RA, Sigma-Aldrich), 1μM of Smoothened-Agonist (SAG, DNSK International) to induce spinal neural progenitors. After 6 days, the cells were fed daily with N2B27 medium supplemented with 1μM RA, 1μM SAG, 5μM DAPT (DNSK International) and 4μM SU5402 (DNSK International) to generate postmitotic spinal MNs. After an additional 7 days (day 14), the cells were dissociated using TryplE (Thermo Fischer Scientific) with added DNaseI (Worthington) and plated onto plates previously coated with Matrigel (BD Bioscences) at a density of 1 million/well of a 6-well plate. Cells were then fed every other day with Neurobasal media supplemented with Non-Essential Amino Acids, Glutamax, N2, B27 and the neurotrophic factors BDNF, CNTF, GDNF (10ng/mL, R&D systems) and Ascorbic acid (0.2μg/ml). In the first four days after plating, the medium was supplemented with EdU (10μM) to inhibit the growth of remaining proliferating progenitor cells. All cell cultures were maintained at 37°C, 5% CO2 without antibiotics and tested regularly for mycoplasma. All cells utilized in the present study were mycoplasma-free.

### Knockdown experiments

For transient knockdown experiments, pre-designed siRNAs (Ambion Silencer Select or Thermo Fisher Scientific) were transfected using Lipofectamine RNAiMAX (Invitrogen) according to manufacturer guidelines. Briefly, for each well of a 6-well plate, we mixed siRNA with lipofectamine RNAiMAX reagent in 200mL Opti-MEM medium and incubated the solution for 20 min at RT. We added 800mL of cell culture medium, delivering a total of 1000mL per well. For *UPF1*, *HBS1L* and *UPF2* KD, 40pmol of a single siRNA were used per each well of a 6-well plate, 5, 10 and 15 days before cell culture collection. For *TDP-43* KD, 20pmol of each of three siRNAs were used 3, 6, 9 and 12 days before cell culture collection. In addition to siRNA treatment, for *TDP-43* KD experiments, a stable iPSC line containing a doxycycline-inducible shRNA targeting *TDP-43* was generated by infecting iPSCs with a lentivirus vector (SMARTvector Inducible Lentiviral shRNA, *hEF1A* promoter) and by selecting for stable cells for 14 days in puromycin (4μM). After differentiation, cells were treated for 15 days with doxycycline (6 μg/mL) before collection.

To knockdown TDP-43 in the context of *UPF2* and *HBS1L* KD experiment, 2.5 uM of ASO targeting *TDP-43* was delivered 14 and 10 days before cell culture collection.

For every experiment, cells were collected and processed 40 days after starting the differentiation process.

### Cellular cytotoxicity assay

Cellular cytotoxicity assay has been performed by measuring Lactate dehydrogenase (LDH) in the media by CyQUANT LDH Cytotoxicity Assay kit (Thermo Fisher Scientific), following the manufacturer instructions. Briefly, 50µL of Reaction Mixture was added to 50µL medium from neurons treated with scramble or UPF1-targeted siRNA, incubated for 30 minutes at room temperature, then arrested with 50uL of Stop solution. Absorbance at 490 and 680 nm was measured with a Synergy LX Multi-Mode Reader (BioTek). Maximum LDH Activity Controls were added with the Lysis Buffer provided with the kit before performing the assay.

### RNA processing and cDNA library constructions for p-UPF1 RIP-seq

iPSC-derived MNs (∼1-2*10^7^ cells) were cultured for 2 hours in okadaic acid (50 nM) before harvesting. The cells were lysed with hypotonic gentle lysis buffer (10 mM Tris (pH 7.4), 10 mM NaCl, 10 mM EDTA, 0.5% w/w Triton X-100) supplemented with Halt Protease and Phosphatase Inhibitor Cocktail (Thermo Fisher Scientific). The lysates underwent immunoprecipitation using Dynabeads Protein A (Thermo Fisher Scientific) with either anti-p-UPF1 S1116 (MilliporeSigma; 07-1016) or rabbit IgG (Sigma Aldrich; I5006) as a control. Cellular RNAs captured by bead-bound antibody complexes were digested to <100 nt by incubating for 30 min at 4°C with RNase I (1 U/μl; Thermo Fisher Scientific Ambion; AM2294). After extensive washing, bound complexes were eluted using IP Elution Buffer (125mM Tris-HCl [pH 6.8], 4% v/v SDS, 20% v/v glycerol, 10% v/v 2-mercaptoethanol, bromophenol blue), and RNA fragments were purified using TRIzol Reagent (Thermo Fisher Scientific). Electrophoresis in 6M urea–15% polyacrylamide parallel to a DynaMarker Prestain Marker for Small RNA (BioDynamics Laboratory; DM253S) was performed. Fragments in the ∼25−50-nt size range were excised, agitated overnight at 25°C in RNA Extraction Buffer (20 mM Tris, 300mM sodium acetate, 2 mM EDTA, 0.2% v/v SDS), and purified using a Coaster Spin-X Centrifuge Tube Filter (Corning; 8162). Further extraction with TRIzol Reagent and ethanol precipitation concentrated the RNA. Control IPs using rabbit IgG were performed, and size-matched Input control samples before IP were prepared by incubating cell lysate for 30 min at 4°C with RNase I in parallel.

The cDNA library is prepared as described(Kurosaki et al., 2021a; Kurosaki et al., 2022). Briefly, to generate libraries for p-UPF1 RIP-seq, purified ∼25−50-nt RNA fragments underwent treatment with recombinant shrimp alkaline phosphatase (New England Biolabs; M0371) to remove 5’-and 3’-phosphates. The fragments were re-phosphorylated at their 5’-hydroxyl groups using T4 polynucleotide kinase (New England Biolabs; M0201S) and purified using the miRNeasy Mini Kit (Qiagen; 217004). A 5’-adenylated DNA adapter (5’-rAppAGATCGGAAGAGCACACGTCT-NH_2_-3’) was added to 3’-ends using truncated T4 RNA ligase 2 (New England Biolabs; M0242S). After ligation of the 5’-RNA adapter (5’-GUUCAGAGUUCUACAGUCCGACGAUC-3’) using T4 RNA ligase 1 (New England Biolabs; M0204S), the RT primer was annealed to the 3’-adapter. Reverse transcription of the adapter-ligated RNAs using SuperScript III reverse transcriptase (Thermo Fisher Scientific) was followed by PCR amplification using Phusion high-fidelity DNA polymerase (New England Biolabs; M0530S) for 15 cycles. PCR products were purified in 8% polyacrylamide, and cDNA quality and quantity were assessed using the 2100 Agilent Bioanalyzer and qPCR. The cDNAs were then sequenced using the Illumina NextSeq550 System.

### RIP-Seq data processing

For RIP-seq data processing, bcltofastq version 2.19.0 was used to demultiplex raw reads generated from the Illumina base calls. Adapter removal and quality filtering was performed using cutadapt. Illumina adapters were removed (-a AGATCGGAAGAGC) along with N calls (--trim-n) and bases with Phred score < 20 (-q 20). A trimmed length of at least 25bp (-m 25) was required to pass filtering. Trimmed reads were aligned to Gencode V38 (GRCh38.p13) with the following flags: --twopassMode Basic; --outSAMtype BAM SortedByCoordinate; -- outSAMstrandField intronMotif.

### RNA preparation

Cells were harversted in 1mL TRIzol Reagent (Thermo Fisher Scientific) per 1-3 million cells. Samples were added with chloroform (0.2mL), mixed and centrifuged at 4°C for 15 minutes at 12,000xg after 3 minutes of resting at room temperature. The aqueous phase containing the RNA was transferred to a new tube and 5µg of RNase-free Glycogen (Thermo Fisher Scientific) have been added. 0.5mL of isopropanol was added to the aqueous phase, the solution was incubated for 25 minutes at room temperature and centrifuged for 10 minutes at 15,000xg at 4°C.

### RNA sequencing

RNA libraries from the first experiment (n=5) were prepared in two batches: 3 samples were sent to Northwestern sequencing core for QC, library preparation (TruSeq Total RNA-Seq Library Prep) and sequencing using the Illumina HiSeq sequencing (PE100, 18 G of raw data per sample); 2 additional samples were sent to Novogene for QC, library preparation (250∼300 bp insert strand specific library with rRNA removal (Ribo-ZeroTM Magnetic Kit)) and sequencing using the Illumina NovaSeq System (PE150, 18 G of raw data per sample). RNA from the following experiments was sent to Novogene for QC, library preparation (Directional mRNA (poly A enrichment)) and sequencing using the Illumina NovaSeq System (PE150, 18 G of raw data per sample).

### RNA-Seq data processing

Raw reads were trimmed using cutadapt (v3.4) to remove Illumina universal adapter sequences (-a AGATCGGAAGAGC), trailing N bases (--trim-n), and bases with Phred score < 10 (-q 10,10), as well as any reads that were too short after trimming (-m 25). Trimmed reads were aligned to Gencode V38 (GRCh38.p13) using STAR (v.2.7.5a). The following flags were used during alignment: --twopassMode Basic, -- outSAMstrandField intronMotif, --outFilterIntronMotifs RemoveNoncanonical, --alignEndsType EndToEnd, as well as the “ENCODE options” flags provided by the STAR manual. Gene-level counts were quantified using featureCounts (v2.0.1). For transcript-level analyses, trimmed reads were pseudoaligned with salmon quant (v1.9.0), using the flags --validateMappings, --gcBias, --seqBias, and --numGibbsSamples 100.

### Differential expression

Gene-level differential expression analysis was performed in R with DESeq2 (v1.38.3). Lowly expressed genes were filtered out before analysis – in order to be retained, a gene must have had at least 0.2 counts per million reads in at least 4 samples. Differentiation was controlled for as a covariate in both RNA-Seq and RIP-Seq analyses. Transcript level differential expression was performed in R with fishpond (v2.4.1), using the Swish pipeline. Briefly, bootstrapped replicates for each sample were adjusted based on library size using scaleInfReps() then transcripts with low expression were marked and removed using labelKeep(). Differential expression analysis was run using the swish() function – differentiation was specified as a covariate during analysis by providing the argument cov=”diff” to swish.

### RNA stability analysis

REMBRANDTS (REMoving Bias from Rna-seq ANalysis of Differential Transcript Stability) was employed to estimate stability of RNA transcripts. As a first step, constitutive exonic and intronic regions were determined for each gene. To determine constitutive exon regions, makeTxDbFromGFF from GenomicFeatures was used to create a TxDb object using transcript annotations from the Gencode V38 GTF. Exonic regions per gene were extracted using exonsBy(txdb, by=”gene”) while intronic regions were extracted per transcript using intronsByTranscript(txdb, use.names=T) and manually aggregated to gene-level. Exonic regions in a gene were considered constitutive if they did not overlap any intronic region for the gene. To determine constitutive intronic regions, transcript locations for each gene were extracted with transcriptsBy(txdb, by=“gene”) and reduced to a single set of coordinates per gene, while exons associated with each gene were extracted with exonsBy(txdb, by=“gene”). Transcript regions were considered constitutive introns if they were not overlapped by any exonic region. Reads were counted for constitutive exonic and intronic regions in R using summarizeOverlaps() from the GenomicAlignments package. For exonic regions, count method was set to “IntersectionStrict” while for intronic regions, the “Union” count method was used. Finally, constitutive intron/exon counts were fed into the REMBRANDTS R script (https://github.com/csglab/REMBRANDTS) and run with default settings. Stability of a gene in control or UPF1 knockout conditions was determined as the mean stability of all samples per condition.

### StringTie analysis

Novel transcripts were assembled for each sample using StringTie (v2.1.3), using the options -G gencode.v38.primary_assembly.annotation.gtf --rf --conservative and default settings otherwise. Transcriptome builds were merged into a unified set of transcripts using StringTie’s merge function along with the same flags. Gffread was used to extract the nucleotide sequence for each transcript. This set of transcripts was used to generate a new decoy-aware Salmon index using the human genome as decoy sequences. Samples were re-quantified against this new transcriptome using the same Salmon protocol as above.

### Transcript feature analysis

ORFs in newly assembled transcripts were predicted using IsoformSwitchAnalyzeR (v2.01.07). Briefly, the merged StringTie.gtf file was loaded using importGTF(). ORFs in known genes were added from the Gencode V38 annotations using addORFfromGTF(), while ORFs from new transcripts were predicted with analyzeNovelIsoformORF(). PTCs were predicted by IsoformSwitchAnalyzeR based on structure of each transcript – transcripts were predicted to have a PTC if the ORF stop codon was >50bp upstream of the final exon-exon junction. Additionally, a manual PTC prediction was implemented in R by introducing cryptic exons into annotated transcript sequences and determining new ORF stop codon locations relative to the final exon-exon junction.

5’ UTR locations and sequences for each protein-coding transcript (i.e. transcripts with a known/predicted ORF) were extracted using ORF annotations. Presence of ORFs in 5’ UTR sequences was evaluated with the findORFs() function from ORFik. A transcript was annotated as having a uORF if the 5’ UTR contained an ATG-to-STOP ORF of at least 10aa. Locations of Alu repeats in the genome were downloaded from RepeatMasker through the UCSC Genome Browser and were intersected with 3’ UTR locations using findOverlaps from the GenomicRanges R package (v.1.50.2) to determine Alu repeat presence. UTR sequences were input into viennaRNA’s RNAfold (v.2.5.0) package to determine UTR structuredness.

### Splicing analysis

Differences in splicing were detected using MAJIQ (v2.4.dev102+g2cae150). Briefly, all samples were input to majiq build along with the flags --min-intronic-cov 1 and –simplify. Majiq deltapsi was employed to quantify splice differences between treatments. Splice events were determined to be significantly different between conditions if dPSI was greater than 0.1 and the probability changing was greater than 0.95. Results tables were extracted using VOILA view with flags --threshold 0.1 and --probability-threshold 0.95 and parsed using custom R scripts.

### Alternate PolyA Usage Analysis

Aligned bam files were converted into bedgraph files using bedtools (v2.30.0-48-g868a9a2-dirty) with the command genomeCoverageBed -bg -ibam <BAMFILE> -split. Basic GencodeV38 annotations (wgEncodeGencodeBasicV38) were downloaded from the UCSC Genome Browser in .bed format and converted into a suitable format for analysis using the DaPars_Extract_Anno.py packaged with DaPars (v 1.0.0). DaPars_main.py was used to run the main analysis using default coverage, FDR, PDUI, and fold change cutoffs.

### Gene ontology and network analysis

The Gene Ontology (GO) analysis was performed using the WebGestalt online tool (Wang et al., 2017). R was used to visualize the GO enrichment. STRING database (Szklarczyk et al., 2021) was used for the visualization of protein functional association. For UPF1 interactions, custom visualizations were created by downloading STRING protein-protein interactions in Cytoscape (v3.9.1) using an interaction confidence cutoff of 0.8. Clustering was adjusted using the organic layout format. After removing unconnected nodes, GO term enrichment was performed within Cytoscape using the stringApp module.

### GSEA

GSEA was performed using fgsea R package (v1.16.0), with Minimal size of a gene set to test=5 and Maximal size of a gene set to test=1000. Wald statistic values (stats) from DESeq2 values from differential expression were used as ranking values for the genes.

### RT-qPCR analysis

The pellet was washed with 1mL of 75% Ethanol and resuspended in DEPC-treated RNase free water (Thermo Fisher Scientific). 0.5 to 1.5µg of RNA was used to generate cDNA using SuperScript IV First-Strand Synthesis System (Thermo Fisher Scientific). Real-time qPCR reactions were prepared according to manufacturer guidelines for PowerUp SYBR Green Master Mix (Thermo Fisher Scientific) on the CFX system (Bio-Rad). To assess gene expression, the Ct value of the housekeeping gene was subtracted from the Ct value of the gene of interest, resulting in the determination of the ΔCt. To assess the cryptic exon level over the total/wild type transcript level, Ct value of the total/wild type transcript was subtracted from the Ct value of the cryptic exon. To assess the relative mature/pre mRNA level, Ct value of the pre-mRNA was subtracted from the Ct value of the mature mRNA. The relative gene expression was calculated as 2^∧^ΔCt (ΔΔCt) and normalized to the control sample.

### Protein processing and Western Blot analysis

Cells were lysed directly into the well and harvested in RIPA buffer (Tris pH 7.4 50mM, NaCl 150mM, Sodium Deoxycholate 0.5%, Triton X-100 1%, SDS 0.2%, EDTA 1mM, protease and phosphatase inhibitors (Millipore)), sonicated for 3×3s, 65V output (QSonica, LLC), centrifuged at 15,000xg for 15min, then quantified for their protein concentration with the Pierce BCA Protein Assay Kit. 20mg of proteins for each sample was mixed with Laemmli SDS loading buffer with added β-Mercaptoethanol and boiled for 5 min at 95°C. Proteins were separated by SDS-PAGE (1-1.5h under 100V constant voltage). In order to separate LC3(II) and LC3(I) isoform, Any kD Mini-PROTEAN TGX Stain-Free Protein Gels (BioRad) were employed. 4–20% Mini-PROTEAN Stain-Free Gels (BioRad) were utilized to study the rest of the targets. Proteins were electro-transferred to nitrocellulose membrane (Bio-Rad) (1h under 100V constant voltage at 4°C). ChemiDoc MP imaging system (Biorad) was used to image the total proteins transferred to the membrane, used as total protein load to normalize the abundance of protein target. After the transfer, the membrane was blocked at room temperature for 2 hours in 5% non-fat milk in TBS and incubated at 4°C overnight with antibodies targeting the protein of interest, diluted in 5% non-fat milk in TBS. The membrane underwent 3×10min washes with TBS-Tween 0.1% and incubated with their corresponding secondary HRP-conjugated antibodies (1:5000, LI-COR Biotechnology) for 1h at room temperature. After 3×10min additional washes with TBS-Tween 0.1%, protein signals were detected by the ChemiDoc MP imaging system using the SuperSignal West Pico PLUS Chemiluminescent Substrate or the SuperSignal West Femto PLUS Chemiluminescent Substrate (Thermo Fisher Scientific). Fiji software (ImageJ, NIH Image) was utilized to perform densitometry analysis of the bands.

### Immunocytochemistry, image acquisition and analysis

Motor neurons were dissociated after differentiation and plated on top of 1.5mm glass coverslips preplated with mouse glia monolayer. Cells were fixed at day 40 with 4%PFA in PBS for 10 min at room temperature. Cells were washed with PBS and permeabilized/blocked at room temperature for 1h with PBS with 0.25% Triton-X and 10% normal donkey serum (Jackson ImmunoResearch). Cells were incubated overnight at 4°C with primary antibody in PBS with 0.1% Triton-X and 2% normal donkey serum, washed 3×10min with PBS with 0.1% Triton-X, and incubated at room temperature for 1h with appropriate secondary antibodies conjugated to BV421, Alexa488, Alexa568 and Alexa647 (1:5000 in PBS with 0.1% Triton-X and 2% normal donkey serum). The coverslips were mounted on glass slides using Fluoromount-G (Southern Biotech). Image acquisition utilized a Nikon A1R+ laser scanning confocal microscope (Northwestern University Center for Advanced Microscopy). Images were captured with consistent exposure times and processed using uniform settings. Images were acquired as Z-stack at 0.3mm intervals, and individual planes were projected to generate maximum intensity images. The NIS-Elements Advanced Research v5 software (Nikon software, Northwestern University Center for Advanced Microscopy) was used to calculate the mean signal intensity of TDP-43, pUPF1, and UPF1 in the regions of interest. 14 to 26 cells per condition were measured for each individual differentiation (n=4). The mean intensity across all the cells in each differentiation was calculated and these valuer were used to run statistic (t.test).

### iCLIP processing and analysis

iCLIP data from HEK293 cells of TDP-43 binding was downloaded from ArrayExpress using accession ID E-MTAB-4733. Raw reads were trimmed using cutadapt, removing adapters from the 3’ end (-a AGATCGGAAGAGC) and 9bp library barcodes from the 5’ end (-u 9), as well as N base calls (--trim-n) and low quality bases (-q 10). Trimmed reads were required to be at least 25bp (-m 25) in order to be retained. Trimmed reads were aligned to Gencode V38 (GRCh38.p13) using ENCODE options from the STAR manual, as well as --twopassMode Basic; --alignEndsType Extend5pOfRead1; --outSAMstrandField intronMotif. Aligned samples were merged using samtools and peak calling was performed on the merged file using PureCLIP (v1.3.1). Peaks were annotated using ChIPSeeker. clip peaks were intersected with APA transcripts using findOverlaps.

### NYGC RNA-Seq data analysis

RNA-Seq data from ALS patients were downloaded from GEO accession GSE137810. Additional patient RNA-Seq data was obtained from Target ALS. Here, patients were included only if they a diagnosis of ALS with no FTD, Alzheimer’s, or other dementia diagnoses. Furthermore, only ALS patients presumed to have TDP-43 pathology (i.e. C9orf72 and sporadic patients) were included – SOD1 and FUS patients were excluded. Additionally, samples with RIN < 5 were excluded, as well as samples missing clinical metadata (e.g. age at death and disease duration). Samples were trimmed using cutadapt with the same flags as prior RNA-Seq samples (-a AGATCGGAAGAGC; --trim-n; -q 10,10; -m 25) and aligned to Gencode V38 using STAR, also with the same flags (--twopassMode Basic; --outSAMstrandField intronMotif; --outFilterIntronMotifs RemoveNoncanonical; --alignEndsType EndToEnd; “ENCODE options” flags from the STAR manual). STMN2 missplicing proportion per sample was determined by counting reads that spanned the STMN2 exon1-exon2 junction and reads that spanned the STMN2 exon1-cryptic exon junction, then calculating (# exon1-CE reads / (# exon1-CE reads + # exon1-exon2 reads).

### Statistical analysis

R (version 4.0.5) and GraphPad Prism (version 9.1.2.226) were used for quantification and statistical analyses.

---

**Supplementary Figure 1.**
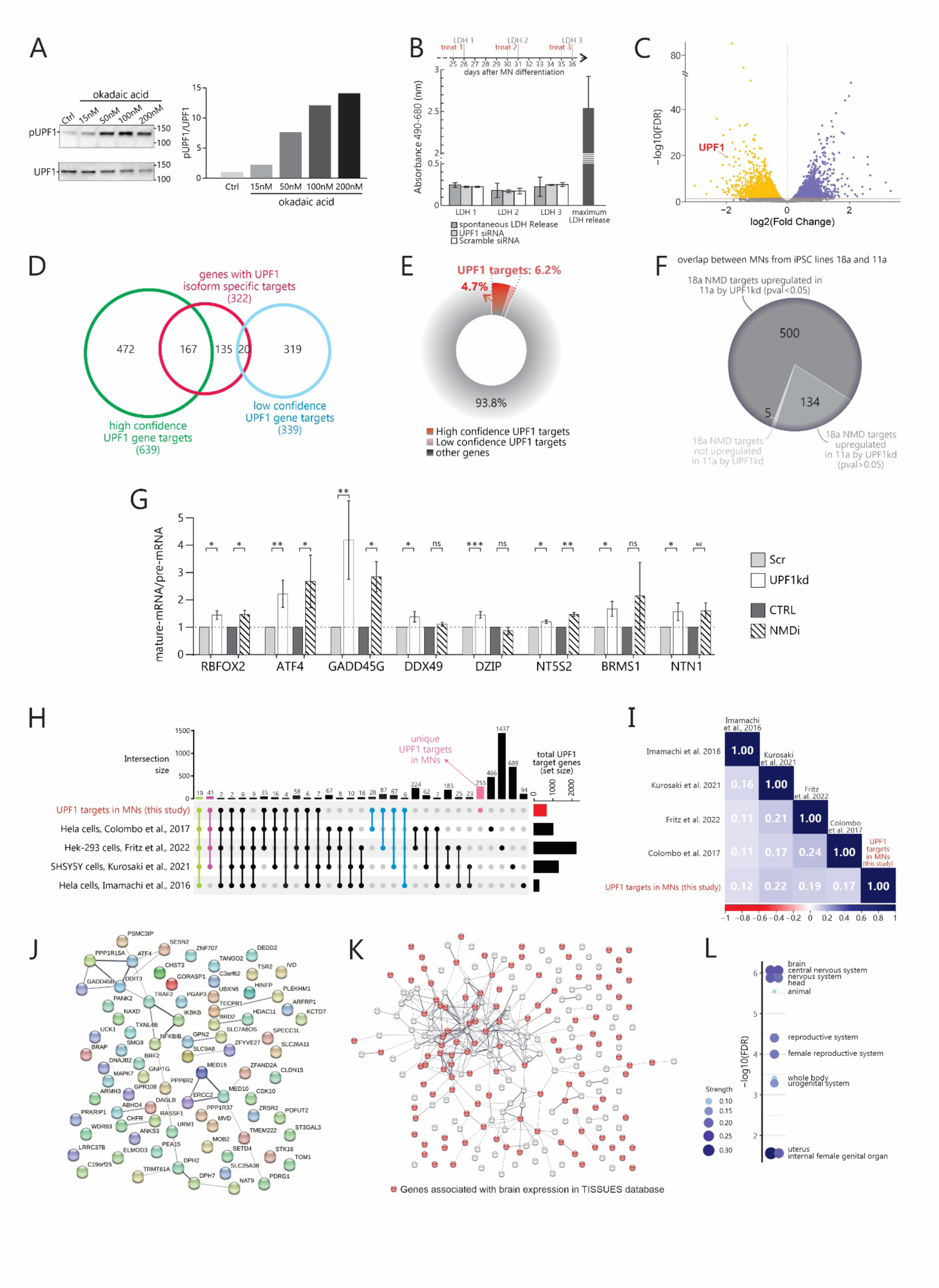
**A)** Left: WB image of pUPF1 and total UPF1 levels in iPSC-derived hMNs untreated or treated for 2 h with different doses of okadaic acid. Right: bar plot showing the pUPF1/total UPF protein ratio in iPSC-derived hMNs untreated or treated for 2 h with different doses of okadaic acid. **B)** Bar plot showing ratio of absorbance at 490 and 680 nm to quantify LDH release following treatment with scramble or UP1 siRNA (n=3 independent differentiations). **C)** Volcano plot displaying differential gene expression analysis of UPF1 KD hMNs. The dot representing *UPF1* gene is highlighted. **D)** Venn diagram showing the overlap of genes with isoform-specific targets (red) with high-confidence (green) and low confidence (light blue) UPF1-target from gene level analysis. **E)** Donut chart showing the percentage of genes (protein-coding and not protein coding) targeted by UPF1 with high and low confidence relative to the total of the genes expressed by hMNs. **F)** Pie chart showing the proportion of UPF1 target genes identified in iPSC line 18a that appeared to be upregulated in iPSC line 11a upon TDP-43 KD in hMNs. **G)** Bar plot showing the fold change of the pre-mRNA/mature mRNA ratio of selected UPF1-target genes in hMNs treated with scramble or UPF1 siRNA and in control and NMDi treated hMNs, analyzed by qPCR (3 to 6 independent differentiations, paired t.test, *p<0.05; **p<0.01; ***p<0.001; ##p<0.001 if t.test performed on technical replicates from 2 independent differentiations). **H)** Upset plot showing all the intersections of UPF1-targets identified in the present study and NMD targets identified in previous studies. In pink are indicated the number of UPF1-targets identified uniquely in the present study. **I)** Correlation plot of UPF1-targets identified in the present study and NMD targets identified in previous studies. **J)** Functional and physical association graph of proteins identified as UPF1 targets in 4 different studies (this study; Colombo et al., 2017(Colombo et al., 2017); Fritz et al., 2022(Fritz et al., 2022); Kurosaki et al., 2021(Kurosaki et al., 2021a)). Line thickness indicates the strength of data support. **K)** Functional and physical association graph of proteins identified as UPF1 targets uniquely in the present study. Line thickness indicates the strength of data support. Red dots represent genes associated with brain expression in TISSUES database. **L)** Bubble plot showing the enrichment analysis of the UPF1 targets uniquely identified in the present study in TISSUES database. The size of the circle is proportional with the strength of the enrichment.

**Supplementary Figure 2.**
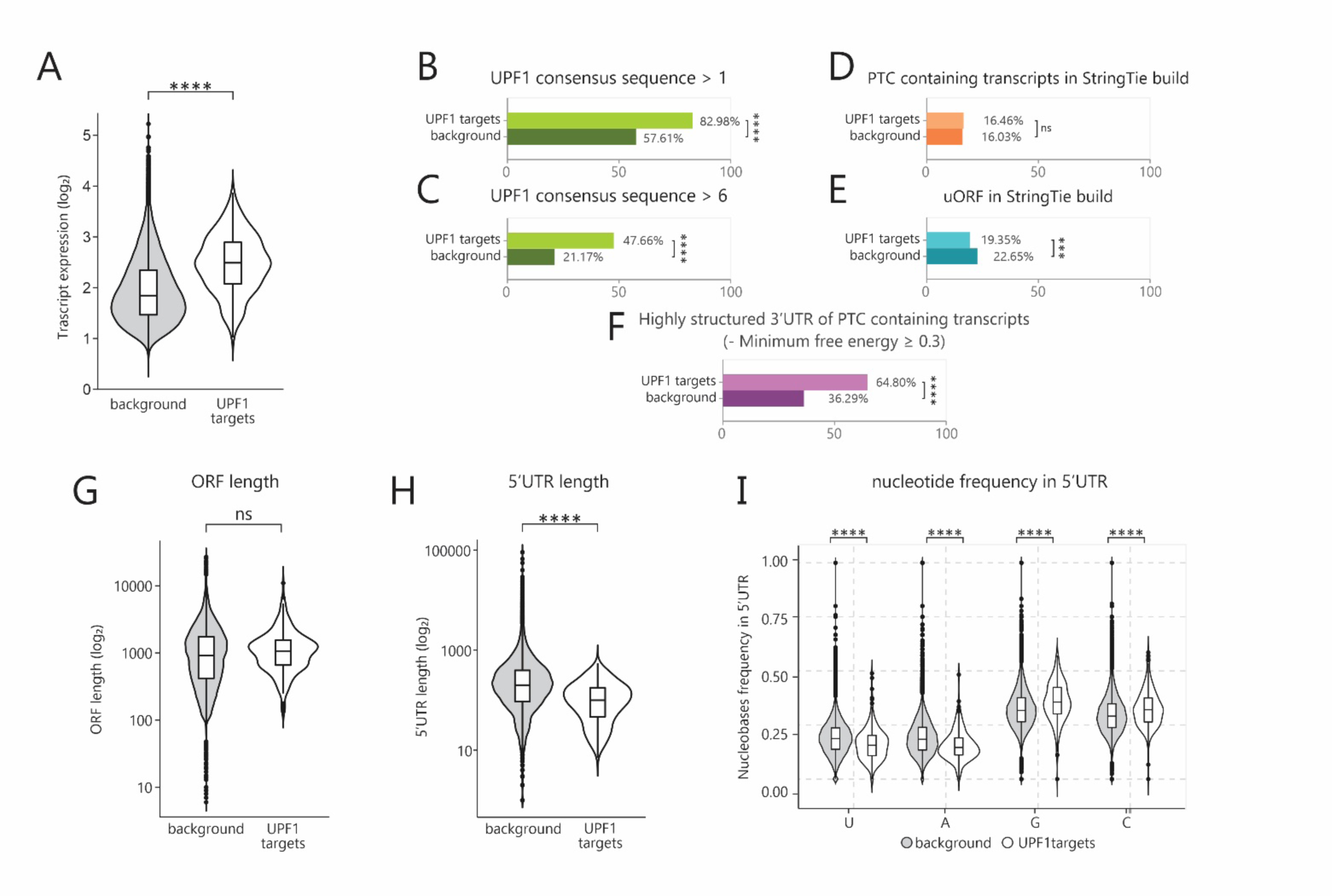
**A)** Violin plot showing expression of transcripts identified as UPF1-targets in hMNs and non-UPF1 targets (t.test, p<0.0001). **B)** Bar plot showing the percentage of UPF1-target transcripts and non-UPF1 targets displaying 2 or more UPF1 binding sequence in the 3’UTR (chi-squared test, p<0.0001). **C)** Bar plot showing the percentage of UPF1-target transcripts and non-UPF1 targets displaying 6 or more UPF1 binding sequence in the 3’UTR (chi-squared test, p<0.0001). **D)** Bar plot showing the percentage of UPF1-target transcripts and non-UPF1 targets harboring a premature stop codon after aligning the RNA-seq data to a custom-build genome including not annotated isoforms (chi-squared test, not significant). **E)** Bar plot showing the percentage of UPF1-target transcripts and non-UPF1 targets presenting upstream open reading frames after aligning the RNA-seq data to a custom-build genome including not annotated isoforms (chi-squared test, p<0.001). **F)** Bar plot showing the percentage of transcripts with highly structured 3’UTR (minimum free energy≥.03) among PTC-containing UPF1-targets, and transcripts with PTCs that are not targeted by UPF1 in our analysis (chi-squared test, p<0.0001). **G)** Violin plot showing the length of ORF of transcripts identified as UPF1-targets in hMNs and non-UPF1 targets (not significant). **H)** Violin plot showing the length of 5’UTR of transcripts identified as UPF1-targets in hMNs and non-UPF1 targets (t.test, p<0.0001). **I)** Violin plots showing the frequency of each nucleotide in the 5’UTR of transcripts identified as UPF1-targets in hMNs and non-UPF1 targets (t.test, p<0.0001).

**Supplementary Figure 3.**
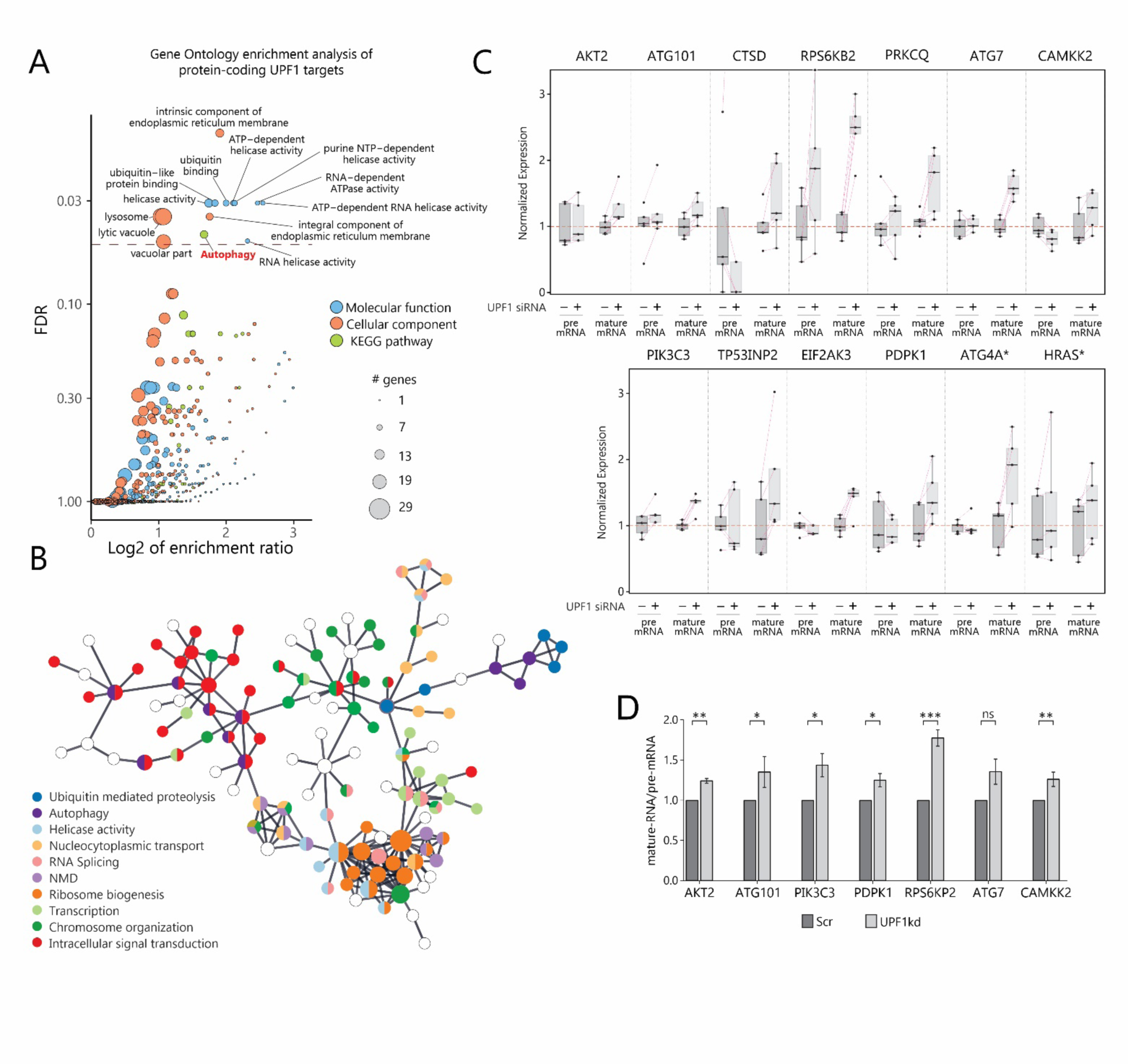
**A)** GO Enrichment analysis (WebGestalt web tool) of the protein-coding UPF1 targets genes. The dashed line indicates the FDR value of 0.05. The size of the circles is proportional to the number of genes of indicated categories. In blue are highlighted the groups associated with the Molecular Function dataset, in orange those with the Cellular Component dataset and in green those with the KEGG pathway. **B)** Functional interconnection analyses of the protein-coding UPF1 targets genes. Colors represent specific annotations as indicated. Not connected nodes or small clusters are not shown. **C)** Boxplots showing pre-mRNA and mature mRNA modulation of autophagy genes after treatment with siRNA targeting *UPF1*. Genes marked with an asterisk after the name indicated that UPF1 targets a PTC-containing isoform. Pink dotted lines connect samples from the same differentiation (n=5 independent differentiations). **D)** Bar plot showing the fold change of the pre-mRNA/mature mRNA ratio of selected genes involved in the Autophagy pathway in MNs treated with scramble or UPF1 siRNA, analyzed by qPCR (3 to 6 independent differentiations, paired t.test, *p<0.05, **p<0.001, ***p<0.0001, ns=not significant).

**Supplementary Figure 4.**
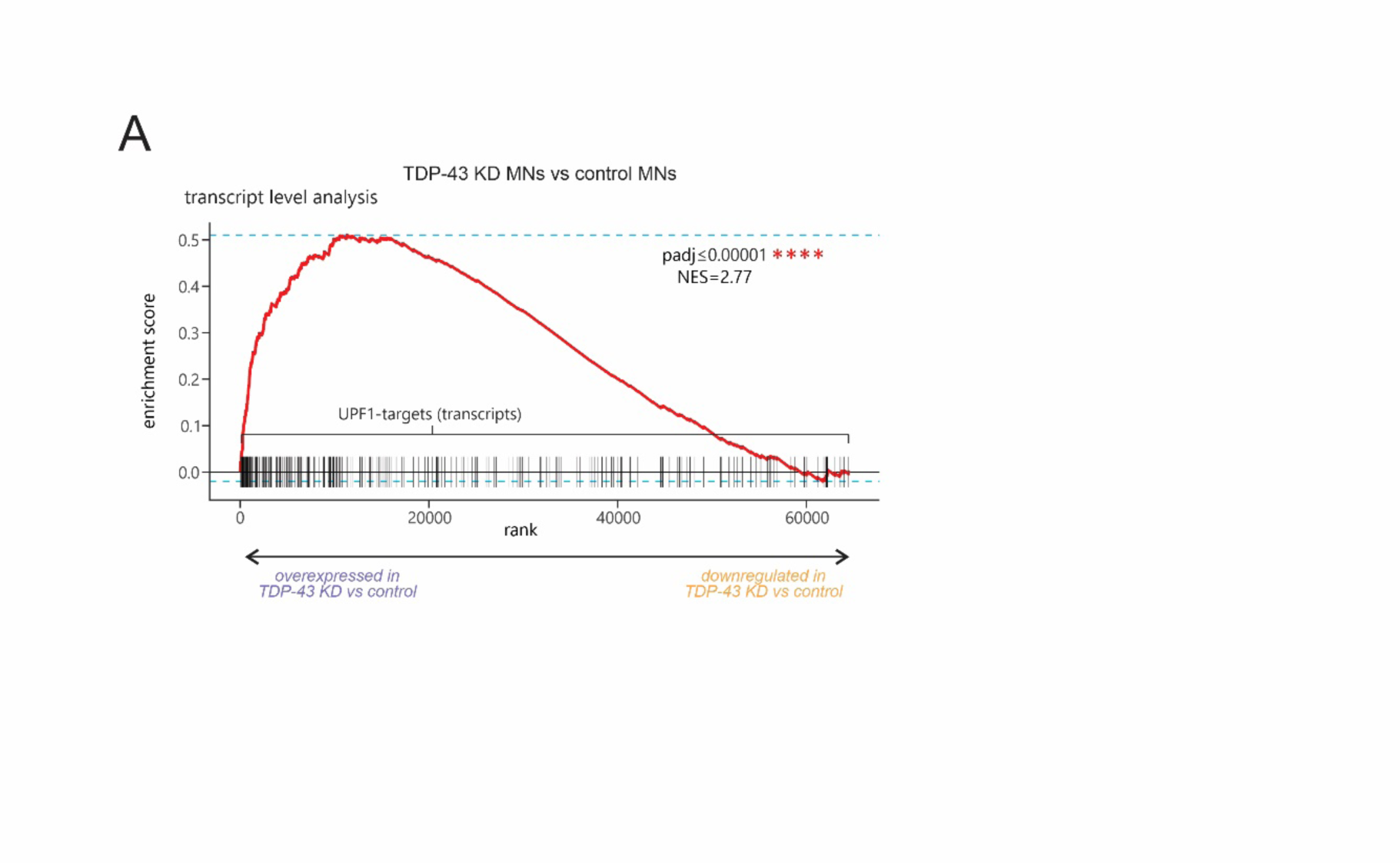
**A)** Enrichment plot from GSEA of TDP-43 KD and scramble siRNA-treated hMNs at transcript level. The plot contains the profile of the running enrichment score (ES) and positions of transcript set elements on the rank ordered list in GSEA. The transcript set includes hMNs UPF1-target transcripts. (NES=Normalized Enrichment Score).

**Supplementary Figure 5.**
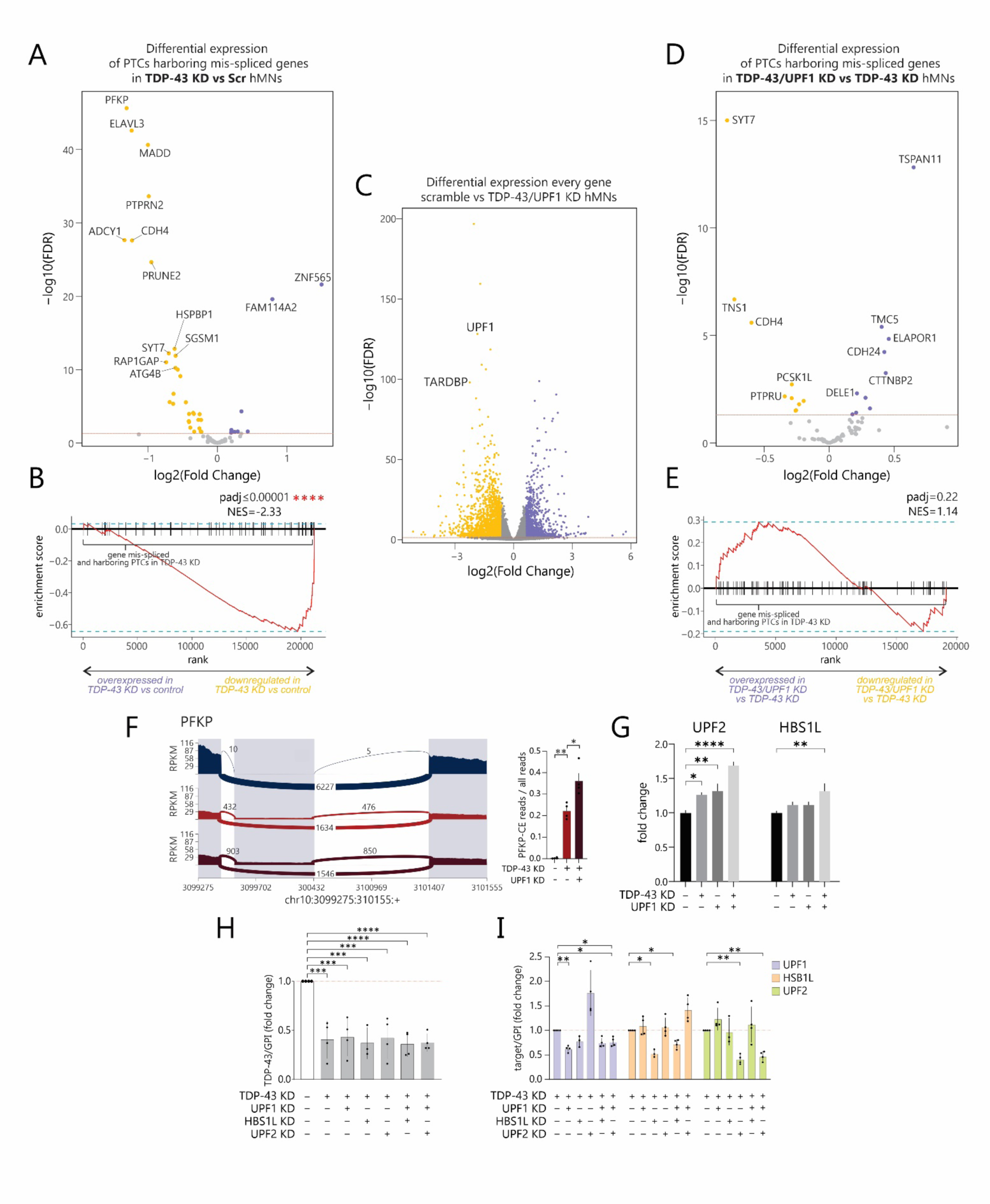
**A)** Volcano plot showing differential expression analysis of TDP-43 splicing targets harboring PTCs in TDP-43-KD and scramble siRNA-treated iPSC-derived hMNs (n=4 biological replicates from 2 individual differentiations). **B)** Enrichment plot from GSEA of TDP-43 KD and scramble siRNA-treated hMNs. The plot contains the profile of the running enrichment score (ES) and positions of gene set elements on the rank ordered list in GSEA. The gene set includes TDP-43 splicing targets genes harboring PTCs. (NES=Normalized Enrichment Score). **C)** Volcano plot displaying differential gene expression analysis of TDP-43/UPF1 KD hMNs. The dots representing *UPF1* and *TARDBP* gene are highlighted. **D)** Volcano plot showing differential expression analysis of TDP-43 splicing targets harboring PTCs between TDP-43/UPF1 KD and TDP-43 KD iPSC-derived hMNs (n=4 biological replicates from 2 individual differentiations). **E)** Enrichment plot from GSEA of TDP-43/UPF1 KD and TDP-43 KD iPSC-derived hMNs. The plot contains the profile of the running enrichment score (ES) and positions of gene set elements on the rank ordered list in GSEA. The gene set includes TDP-43 splicing targets genes harboring PTCs. (NES=Normalized Enrichment Score). **F)** Left: Sashimi plots showing mean cryptic event occurrence in *FPKP* in scramble siRNA treated, TDP-43 KD and TDP-43/UPF1 KD hMNs. Right: Bar plot showing quantification of CE-containing reads/total reads ratio of *FPKP* from RNA-seq of scramble siRNA treated, TDP-43 KD and TDP-43/UPF1 KD conditions (n=4 biological replicates from 2 individual differentiations; t.test, *p<0.05; **p<0.01). **G)** Bar plot showing the expression level of *UPF2* and *HBS1L* upon UPF1, TDP-43 and TDP-43/UPF1 KD conditions over scramble-siRNA treated hMNs by RNA-seq (*FDR< 0.05, **FDR<0.01, **** FDR<0.0001). **H)** Bar plot showing the level of *TARDBP* in TDP-43-KD hMNs treated with siRNA targeting UPF1, HBS1L and UPF2 as indicated in the figure, relative to a scramble ASO control. Expression values are normalized to the housekeeping GPI gene level. (n=4 biological replicates from 2 individual differentiations, One-way ANOVA followed by Dunnett multiple comparison test, ***p<0.001, ****p<0.0001). **I)** Bar plot showing the level of UPF1 (light purple), HBS1L (light orange) and UPF2 (light green) in TDP-43-KD hMNs treated with siRNA targeting UPF1, HBS1L and UPF2 as indicated in the figure. Expression values are normalized to the housekeeping GPI gene level. (n=4 biological replicates from 2 individual differentiations, One-way ANOVA followed by Dunnett multiple comparison test, *p<0.05, **p<0.01).

**Supplementary Figure 6.**
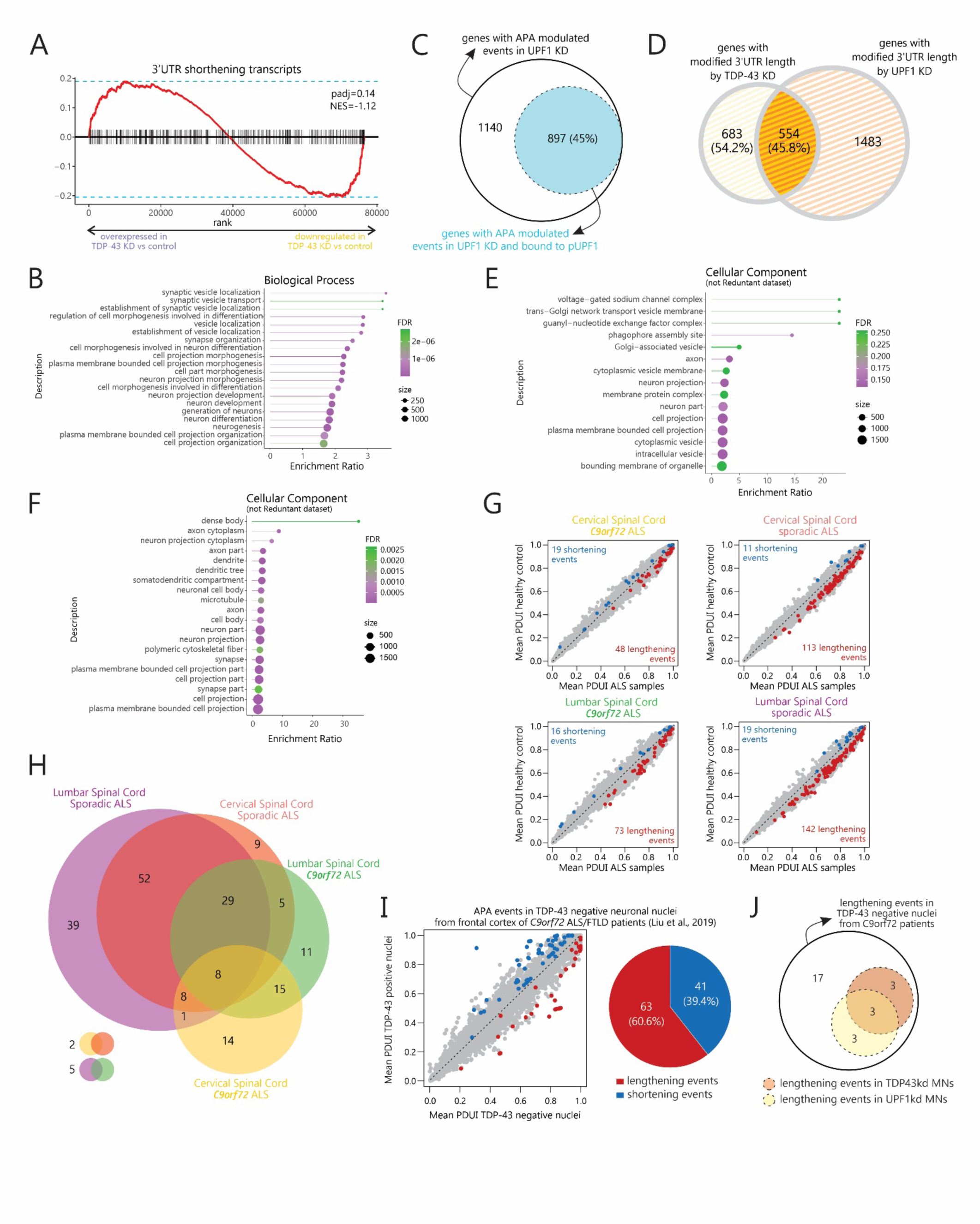
**A)** Enrichment plot from GSEA of TDP-43 KD and scramble siRNA-treated hMNs. The plot contains the profile of the running enrichment score (ES) and positions of gene set elements on the rank ordered list in GSEA. The gene set includes hMNs genes displaying 3’UTR shortening events upon TDP-43 KD. (NES=Normalized Enrichment Score). **B)** GO Enrichment analysis (Biological Process Dataset) of the genes undergoing APA by TDP-43 KD. Only the most significant 20 categories are shown. The color indicates the FDR score, the size of the circles is proportional to the number of genes of indicated categories. **C)** Euler plot showing APA genes in UPF1 KD hMNs that are significantly enriched in pUPF1 RIP-seq. **D)** Venn diagram showing the intersection of APA genes upon TDP-43 KD and UPF1 KD hMNs. Percentages indicate the proportion of TDP-43 genes overlapping or not overlapping UPF1 events. **E)** GO Enrichment analysis (Cellular Component not redundant Dataset) of the genes undergoing APA resulting in shorter 3’UTR by both TDP-43 KD and UPF1 KD. The color indicates the FDR score, the size of the circles is proportional to the number of genes of indicated categories. **F)** GO Enrichment analysis (Cellular Component not redundant Dataset) of the genes undergoing APA resulting in longer 3’UTR by both TDP-43 KD and UPF1 KD. Only the most significant 20 categories are shown. The color indicates the FDR score, the size of the circles is proportional to the number of genes of indicated categories. **G)** Dot plots showing the APA events occurring in human cervical (top) and lumbar (bottom) spinal cord of ALS patients segregated among *C9ORF72* ALS (left) and sporadic ALS (right) patients. **H)** Venn diagram showing the intersection of APA events occurring in human cervical (top) and lumbar (bottom) spinal cord of ALS patients segregated among *C9ORF72* ALS (left) and sporadic ALS (right) patients. Red and blue represent respectively lengthening and shortening statistically significant events. **I)** Left: Dot plots showing the APA events occurring in TDP-43 negative neuronal nuclei isolated from the frontal cortex of *C9ORF72* ALS patients(Liu et al., 2019). Red and blue represent respectively lengthening and shortening statistically significant events. Right: Pie charts showing the proportion of lengthening (red) and shortening (blue) events occurring in TDP-43 negative neuronal nuclei. **J)** Euler plot showing the APA events occurring in TDP-43 negative neuronal nuclei that we observed by knocking down TDP-43 (yellow circle) or UPF1 (orange circle) in iPSC-derived hMNs.

